# Bioprinting Taurine-Incorporated Gelatin Methacrylate Hydrogels for Enhanced Muscle Tissue Regeneration

**DOI:** 10.1101/2025.04.25.650478

**Authors:** Nima Tabatabaei Rezaei, Kartikeya Dixit, Sangeeta Shrivas, Hitendra Kumar, Keekyoung Kim

## Abstract

Skeletal muscle diseases like myopathies and muscular dystrophies present significant clinical challenges with few effective treatments. To better understand disease mechanisms and accelerate therapy development, robust in vitro muscle models are needed. Extrusion- and light-based bioprinting offer precise fabrication of tissue-like constructs, but creating bioinks that support muscle cell function remains difficult. Here, we report a novel bioink in which taurine is first methacrylated—substituting its –NH_2_ groups to afford taurine methacrylate (TMA)—enabling covalent integration into gelatin methacryloyl (GelMA) networks. We systematically compared GelMA-Taur (physical blend) versus GelMA-TMA (covalent) hydrogels, assessing mechanical stiffness, swelling behavior, and photocrosslinking kinetics. Incorporating TMA yielded tighter crosslinking control, minimized overcure in DLP-printed features, and improved shape fidelity. SEM revealed finer pore structures and homogeneous TMA distribution, and release assays confirmed prolonged TMA retention compared to rapidly leaching taurine. Photopatterning and 3D printing of complex geometries demonstrated excellent printability of the GelMA-TMA bioink. Finally, C2C12 myoblasts encapsulated in GelMA-TMA scaffolds exhibited accelerated early differentiation, higher myosin heavy chain expression, and more extensive myotube formation than controls. Together, these results establish GelMA-TMA as a printable, mechanically tunable, and biologically active platform for engineering skeletal muscle tissue in disease modeling and regenerative applications.

## 1. Introduction

Skeletal muscle is the predominant tissue type in an adult body, comprising around 40-45% of the body mass [1,2]. It is made up of bundles of contractile, multinucleated muscle fibers that produce the forces needed for essential physiological activities like walking, chewing, and eye movement [3,4]. Skeletal muscles are intrinsically capable of repairing minor injuries (e.g., strains); however, effective regeneration is highly orchestrated through multiple cellular responses [5,6]. Muscle repair mechanism resident multipotent stem cells, called myosatellite cells, are activated and rapidly proliferate and differentiate into myoblasts [6]. These myoblasts then fuse to create new myofibers, regenerating the structural and functional integrity of the muscle tissue. Except for those cases of trauma, it has been found that when injury exceeds 20% of the original muscle mass, innate regenerative capacity is insufficient [7]. This severe damage results in fibrotic scar tissue deposition, which misarranges the native biological milieu and microarchitecture of the muscle [6]. This leads to considerable functional impairment. The clinical gold standard for reconstructing large-scale muscle defects is autologous tissue transplantation, such as muscle flaps [8]. Although this is effective in some cases, it has its limitations, including a scarcity of donor tissue, functional loss at the donor site, and morbidity from tissue harvesting [8]. Skeletal muscle tissue engineering (SMTE) can be used as a potential alternative to overcome these challenges [9].

SMTE provides an innovative approach for limited regeneration of injured skeletal muscle through a combination of cells, advanced biomaterials, and bioengineering approaches. Such an approach aims to restore the structural muscle and recover its function, addressing the limitations of current clinical treatments. Such strategies can broadly be categorized into two approaches: (1) tissue engineering of complex muscle constructs *in vitro* for implantation and (2) tissue-like scaffolds development enhancing muscle regeneration *in vivo* [10]. Both these approaches are based on a combination of scaffolds, cells, and molecular signaling, but differ in their relative emphasis. The former focuses on artificial muscle tissue biofabrication, while the latter aims to support muscle tissue formation and regeneration *in vivo*. Due to the dynamic nature of skeletal muscle, successful SMTE strategies require the use of biomaterials compatible with muscle progenitors and the design of appropriate 3D architectures to support functional regeneration [11]. There are various approaches for biofabrication scaffolds in tissue engineering, with three-dimensional (3D) bioprinting emerging as a highly promising method [12]. This advanced technique is widely employed for regenerating severely damaged muscle tissue. By combining different types of cells with biomaterials, 3D bioprinting enables the fabrication of structures that support cell attachment, growth, and differentiation [13]. These constructs facilitate the formation of mature tissue with high cell viability, making them suitable for a range of biomedical applications. Numerous naturally derived biomaterials have been explored in literature for bioprinting muscle tissue [14] For instance, Haas et al. [15] engineered biomimetic collagen-gelatin–laminin sponges cross-linked with genipin that mimic native muscle (ECM) extracellular matrix architecture and mechanical cues, promoting vascularized, innervated muscle regeneration and improved contractile force. Additionally, Gupta et al. [16] provided a systematic evaluation of gelatin hydrogels enzymatically cross-linked, which robustly guided C2C12 and primary chick myoblast alignment, fusion, and sarcomere maturation, highlighting their promise for engineered skeletal muscle models.

Gelatin is one of the most commonly used natural polymers in muscle tissue *in vitro* bioprinting due to its Arg-Gly-Asp (RGD) peptide sequences, which promote cell adhesion [14,17]. Previous studies have demonstrated that gelatin, in combination with laminin and collagen, supports the invasion and survival of C2C12 myoblasts in 3D cultures [15]. It also enhances the secretion of myogenic markers such as MyoD and MyoG and promotes the proliferation of endothelial and satellite cells when used in volumetric muscle loss (VML) defects. However, these beneficial effects are limited to approximately two weeks, after which myofiber restoration is no longer sustained [15]. To address these limitations, the mechanical strength of gelatin has been enhanced by introducing methacryloyl groups onto its backbone, resulting in gelatin methacrylate (GelMA) [18,19]. GelMA is a promising biomaterial for stereolithography (SLA) and digital light processing (DLP) bioprinting due to its tunable mechanical properties, thermosensitivity, and rapid gelation kinetics [20]. To further enhance the mechanophysiological and bioactive properties of GelMA hydrogels, it has been compounded with various additives, including other natural or synthetic polymers [21–24], nanoparticles [25,26], and small molecules [27,28]. These modifications expand its applicability and effectiveness in tissue engineering and regenerative medicine.

Recently, small molecules with antioxidant properties have garnered significant attention in the biomedical field due to their ability to act as reducing agents, hydrogen donors, free radical scavengers, and singlet oxygen quenchers [29–31]. Taurine (2-aminoethanesulfonic acid, Taur) is one such molecule—a sulfur-containing, non-protein, non-essential amino acid with critical physiological, pathological, and biological roles in various tissues and organs of the human body [32–34]. These roles include osmoregulation, antioxidation, and maintaining intracellular homeostasis [35]. Notably, Taur has a biologically specific function in modulating the phosphatidylinositol 3-kinase (PI3K)/Akt signaling pathway, which regulates the cell cycle and supports cell proliferation across various cell types [36]. Its necessity for proper muscle functioning has been well-documented. For instance, studies using Taur transporter knockout mouse models have demonstrated that the absence of Taur leads to impaired muscle conductance velocity without affecting nerve conductance speed, alongside significantly reduced exercise performance and running speeds [37,38]. Furthermore, *in vitro* studies have highlighted Taur ‘s critical role in muscle cells. Overall, by regulating the PI3K/AKT, AKT/FOXO1, JAK2/STAT3 and mTOR/AMPK signal pathways, Taur can increase cell proliferation and protein synthesis [39,40]. Also, Taur has been shown to enhance myoblast differentiation into myotubes via the Taur transporter and the Ca^2+^ signaling pathway, underlining its importance in muscle development and function [41]. These findings collectively emphasize Taur’s potential as a bioactive molecule for therapeutic applications in muscle health and regeneration.

In this work, Taur was functionalized by substituting its —NI_2_ groups with methacrylate groups, enabling covalent bonding between Taur and the GelMA backbone to introduce bioactivity for tissue engineering applications. The incorporation of Taur and TMA into GelMA was followed by an investigation of its mechanical and physiological properties. Furthermore, the effect of covalent bonding between Taur/TMA and GelMA on its retention within the hydrogel structure was examined by studying the release kinetics. In addition, Taur-containing GelMA was 3D printed to assess its potential for fabricating complex bioengineered structures. To evaluate biological performance, C2C12 myoblasts were encapsulated within the modified GelMA scaffolds, and the impact of Taur and TMA on cell proliferation and Myosin Heavy Chain (MHC) secretion was analyzed.

## 2. Materials and Methods

### 2.1 TMA synthesis and bioinks preparation

Taurine methacryloyl (2-methacrylamidoethane-1-sulfonic acid, referred in this study as TMA) was synthesized by reacting Taurine (2-aminoethane-1-sulfonic acid) (Sigma Aldrich, St. Louis, MO, USA) with glycidyl methacrylate (GMA) (Sigma Aldrich, St. Louis, MO, USA). The synthesis was performed using distilled water as the solvent and combinations of different reaction conditions as specified in Table 1. First, 2 g of Taur was dissolved in 20 mL of distilled water. Next, GMA was added dropwise to the Taur solution while maintaining continuous stirring at 500 rpm. The reaction was continued for 24 hours in the dark with continuous stirring at 500 rpm. Next, the resulting solution was added dropwise to 100 mL cold ethanol with continuous stirring at 200-250 rpm. After complete addition of the solution, the resulting mixture was left undisturbed for 10 minutes. Next, the mixture was agitated and transferred to 50 mL tubes and centrifuged to collect the precipitate. The supernatant was carefully removed, and 30 mL of fresh ethanol was added to each tube, followed by vigorous agitation and centrifugation. This process was repeated three times to obtain a filtered precipitate of TMA. Next, the precipitate was resuspended in 20 mL ethanol, poured on a glass petri dish, and dried at 50 °C. The dried powder of TMA was then stored at -20 °C for further usage.

### 2.2 Methacrylation evaluation (NMR)

In order to evaluate the methacrylation process and quantify the degree of substitution of the synthesized TMA, proton nuclear magnetic resonance (^1^H NMR) was used following previously established methods [42,43]. A 5% (w/v) solution of Taur and TMA was prepared by dissolving the sample in 1 mL of deuterium oxide (ThermoFisher, Waltham, Massachusetts, USA). The NMR spectra was recorded using a 600 MHz Bruker Avance III Spectrometer. Methylene protons at around 3.2 ppm were selected as the internal reference. Methacryloyl proton signals at 5.7 ppm, 6.2 ppm, and 1.97 ppm were integrated to determine the degree of substitution [44].

### 2.3 Mechanical and swelling properties evaluation

The mechanical properties of the hybrid hydrogels were assessed by determining the compression modulus, which reveals the microstructural stiffness. To prepare the samples, different concentrations of Taur and TMA were dissolved in 5% GelMA solution containing 0.2% LAP and placed into molds measuring 8 mm in diameter and 8 mm in height, with each well receiving 500 μl of the respective sample, fhe prepolymer solution was then subjected to visible light crosslinking for 3 minutes to ensure uniform crosslinking throughout the entire volume of the samples.

A vertical axis micromechanical testing machine, equipped with a flat-ended rigid cylinder (12.5 mm diameter) as the compression probe, was utilized to record force-displacement plots for the hydrogels during indentation. The hydrogels were subjected to approximately 70% displacement at a speed of 20 mm/min to generate these plots. Using the initial dimensions of the samples and an in-house MATLAB code, the compression modulus was calculated based on the slope within the initial 10% strain region, representing the linear region of the stress-strain curve for 5 replicated for each combination.

Determining the swelling ratio of the crosslinked hydrogel samples is crucial for evaluating their water uptake capacity. Briefly, 250 μl of the hybrid hydrogel solutions were cast into molds with a diameter of 8 mm. These samples were then crosslinked, then immersed in PBS and incubated at 37°C for 24 hours to ensure full hydration. Once fully hydrated, the wreight of each sample in its hydrated state (w_w_) was measured. Subsequently, the hydrated hydrogel samples were frozen at -80°C and lyophilized for two days to determine their dry weight (w_d_). The swelling ratio was calculated using **Equation (1)**, with five replicates for accuracy.

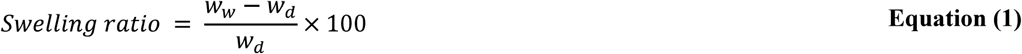

### 2.4 Gels conductivity measurement

The electrical conductivity of the hydrogels was assessed through electrochemical impedance spectroscopy (EIS) using a PalmSens4 device (PalmSens BV, Netherlands). In general, hydrogel samples were prepared with a composition of 5% w/v GelMA and varying concentrations of Taur and TMA at 1%, 5%, and 10%. The photo-crosslink:ing process was set to a fixed duration of 2 minutes for all samples, with the hydrogel dimensions standardized to 24 mm × 24 mm × 2 mm (L×W×H). Additionally, these hydrogel samples were prepared using PBS as the solvent. A custom-made 3D-printed platform was designed to hold the samples between electrodes, which were constructed from glass slides coated with conductive copper tape. After crosslink:ing, each hydrogel sample was carefully wiped to remove any excess liquid before being positioned between the two electrodes. The ends of these electrodes were connected to the EIS station.

Conductivity measurements were performed at room temperature over a frequency range spanning from 0.1 Hz to 1 MHz, with an applied potential set at 10 mV and an amplitude of 1 mV. The data were acquired and subsequently analyzed using PSTrace software (PalmSens BV, Netherlands). The conductivity (σ) of the hydrogels was then calculated according to **Equation (2)**, where (*l*) represents the distance between electrodes, and (*A*) denotes the electrode area in contact with the hydrogel. The resistance (*R*) of the hydrogel samples was determined by fitting the Nyquist plot to a Randles equivalent circuit model.

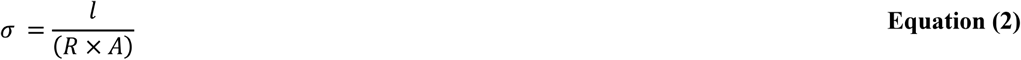

### 2.5 Transparency evaluation

The transparency of the GelMA-Taur/TMA hydrogels was evaluated usmg UV–Vis spectroscopy (Shimadzu UV 2600, United States of America). Measurements were performed with square-bottom cuvettes having a width of 5 mm, spanning a wavelength range of 350 to 950 nm. Prior to the measurements, the hydrogels were prepared by vortexing and then centrifuging at 1500 rpm for 1 minute. This process removed air bubbles and ensured sample uniformity, allowing for accurate transparency assessments.

### 2.6 Photocrosslinking kinetic evaluation

The photocrosslinking kinetics of the GelMA Taur/TMA hydrogels were assessed using an Anton Paar MCR 302e rheometer (Anton Paar, Austria). For measurements, a stainless-steel measuring set having 25 mm parallel plate geometry was used. Prior to initiating the photo-crosslinking process, the hydrogels were carefully prepared as described in previous sections and loaded onto the rheometer to ensure uniform distribution between the parallel plates. The gap between the plates was adjusted to 1 mm maintaining consistent sample thickness, and any excess material was gently removed to avoid edge effects. Initially, the hydrogels were probed to measure their storage modulus (G’) and loss modulus (G”) under oscillatory shear conditions for 1 minute in the absence of irradiation. This step allowed for the determination of the baseline viscoelastic properties of the uncrosslinked hydrogel. During the measurement, the frequency was set at 1 Hz, with a strain amplitude within the linear viscoelastic region (1%) to avoid altering the structure of the sample. Following the baseline measurement, the hydrogel sample was subjected to UV irradiation for 5 minutes from the bottom using a UV light source integrated into the rheometer. During irradiation, the storage and loss moduli were continuously monitored in real-time to capture the dynamic changes in viscoelastic properties as the hydrogel underwent photo-crosslinking.

The photo-crosslinking kinetics were evaluated by analyzing the evolution of G’ and G” over the irradiation period. A sharp increase in G’, accompanied by a plateau, indicated the completion of the crosslinking process and the formation of a stable hydrogel network. The crossover point between G’ and G”, where G’ surpassed G”, was identified as the gelation point, marking the onset of significant network formation. After the 5-minute irradiation period, the UV light source was turned off, and the samples were subjected to a frequency sweep. This additional step was conducted to evaluate the viscoelastic properties of the fully crosslinked hydrogels across a range of frequencies (0.1 to 100 rad/s), providing insights into their stability and mechanical behavior under varying oscillatory conditions. To ensure reproducibility and statistical validity, all experiments were conducted in triplicate under controlled conditions, including a constant temperature of 23 °C.

### 2.7 Scanning electron microscopy (SEM)

To investigate the microstructure of the GelMA-Taur/TMA hydrogels, we utilized scanning electron microscopy (SEM) to see the distribution of the Taur/TMA and its influence on structural changes. Each hydrogel group was prepared by mixing 250 µl of hydrogel precursor at specified concentrations, which were then crosslinked in a cylindrical container under visible light. After crosslinking, the samples were frozen at -80 °C and lyophilized for two days. Following lyophilization, the samples were sputter-coated with gold using a sputtering machine. The coated samples were then imaged using a Phenom ProX benchtop SEM to analyze the microstructure. Furthermore, the pore size of the samples was analyzed by ImageJ based on the SEM images.

### 2.8 Taurine release kinetics

To evaluate the retention of Taur and TMA within the GelMA structure over time, their cumulative release was measured at specific incubation intervals. GelMA-Taur/TMA hydrogels with previously mentioned concentrations were prepared by casting 400 µL of each formulation into cylindrical molds (8 mm in diameter) and photo-crosslinking under visible light for 3 minutes to ensure complete crosslinking. For each group, five samples were placed in 6-well plates, immersed in 3 mL of PBS, and incubated at 37 °C on an orbital mixer at 70 RPM. At predetermined time points, 500 µL of supernatant was extracted from each well for further analysis and stored at –20 °C. To maintain consistent volume, an equal amount (500 µL) of prewarmed fresh PBS was added back to each well after every sampling. After 4 days, all collected samples were analyzed to quantify Taur concentration using a Taur ELISA kit (AFG Scientific) following the manufacturer’s protocol. The cumulative release of Taur was then calculated and expressed as a percentage of the initial loading concentration for each group.

### 2.9 Photo-patterning and 3D printing

To evaluate the printability of the bioink:s, a 450 µm-thick layer of 5G and 5G10TMA was dispensed onto a glass petri dish and crosslink:ed into various patterns using a custom-built DLP printer equipped with a 405 run projector. To compare printing resolution, the line thickness of different sections was measured using ImageJ software.

For the additive manufacturing of thicker structures, a modified Anycubic Photon Mono 4K DLP printer with a 405 run light source was utilized. Various structures were printed with a layer thickness of 100 µm and an exposure time of 15 s per layer. Additionally, the first six bottom layers were crosslinked with a 40 s light exposure to ensure a stable base.

### 2.10 Cell culture and cell-laden hydrogels viability assessment

Immortalized mouse myoblast cell line (C2C12) cells (sourced from ATCC) were cultured in Dulbecco’s modified Eagle medium (DMEM) (Lonza, Basel, Switzerland) enriched with 10% fetal bovine serum (FBS) and 1% penicillin/streptomycin (PEN/ST) incubated in 37°C and 5% CO_2_. The adherent cells were detached using trypsin-EDTA solution (Sigma-Aldrich), centrifuged, and then resuspended in prepolymer hydrogels at a density of 3 × 10^6^ viable cells per mL. Samples were crosslinked in a cylindrical mold with 6 mm diameter and 450 µm thickness using homemade DLP printer.

Cell viability analysis was performed using the LIVE/DEAD assay Kit (Biotium, Fremont, CA, USA). For assessments of cell viability and growth, the cell-laden structures were crosslinked and then cultured in the same growth medium, with media changes every 3 days. On days 3, 5, and 7 of culture, 5mm disk samples were punched out and after three PBS washes, the encapsulated cells were stained for 30 minutes in the dark at room temperature using a PBS solution containing 0.5 µL mL^−1^ CalceinAM and 2 µL mL^−1^ EthD. Subsequently, the samples were washed three more times with PBS to remove any residual reagents. Fluorescence images were captured using an inverted fluorescence microscope, with Calcein AM fluorescence detected in the FITC channel and EthD fluorescence in the TXR channel.

### 2.11 Proliferation and morphological assay

In addition to viability assay, cells were encapsulated within hydrogel scaffolds and cultured for one month to characterize and visualize cell proliferation within GelMA and GelMA-Taur/TMA bioinks. To investigate cell adhesion and morphology, encapsulated samples were punched using a 5 mm surgical punch, and at least three disks from each sample were stained with phalloidin (Cytoskeleton, Denver, CO, USA) for cytoskeleton staining and DAPI (Fluoroshicld™ with DAPI, Sigma-Aldrich) for nuclei staining at 7, 10, 14, 21, and 28 days of culture.

The staining process involved washing the hydrogel samples with PBS three times, fixing them in 4% paraformaldehyde for 90 minutes, permeabilizing the cell membranes with 0.5% Triton X-100 in PBS for 20 minutes, and incubating with 500 μ l of phalloidin 488 stock solution at room temperature for 90 minutes. Following another scries of PBS washes, the samples were transferred to glass slides, treated with DAPI stock solution, covered with a cover slip, and imaged using 10× and 20× objective lenses on an inverted fluorescence microscope (ECHO Revolve) equipped with DAPI and EGFP channels.

A minimum of five images per sample were analyzed using ImageJ software to precisely quantity the size and number of clusters within a given unit area.

### 2.12 Image analysis and myotubes formation quantification

Using custom MATLAB scripts, we tracked and analyzed cell chains. Briefly, each FITC and DAPI image was converted to grayscale, and cell chains were manually annotated by marking their nuclei. We then quantified the number of nuclei per myotube and measured each myotube’s length. For each condition, five images (10× magnification) were analyzed, and those data arc reported.

### 2.13 3D Bioprinting

To assess cell fate within the scaffolds, 450 µm-thick scaffolds of 5G and 5G10TMA, containing 3 × 10^6^ viable cells per mL, were bioprinted using Anycubic Photon Mono 4K DLP printer various patterns were photocrosslinked, followed by the removal of uncrosslinked regions through washing. The 3D-fabricated, cell-laden structures were then cultured in complete culture medium and incubated at 37°C with 5% CO_2_. After seven days of culture, Phalloidin/DAPI staining and immunostaining were performed to analyze cell morphology and myotube formation within the scaffolds.

### 2.14 Immunostaining and confocal microscopy

To visualize myotube formation, immunostaining was performed on fixed, Phalloidin-stained, cell-laden structures. The samples were incubated overnight at 4°C with Anti-Fast Myosin Skeletal Heavy Chain antibody (MY32) (1:500, Abeam, ab51263). Following incubation, the samples were washed three times with PBS and then treated with goat anti-mouse Alexa Fluor 594 (1:1000, Abeam, abl 50120) for 2 hours at room temperature. After an additional three PBS washes, the samples were mounted with DAPI-containing mounting media. Fluorescence imaging was performed using an inverted fluorescence microscope (ECHO Revolve) with DAPI, FITC, and TRX channels. Through image analysis, the MHC-positive index was calculated as the percentage of MHC-positive cells. Additionally, the fusion index was determined as the percentage of MHC-positive cells containing more than two nuclei [45]. A Nikon Eclipse Ti confocal microscope (Nikon, Tokyo, Japan) was used to perform high-resolution 3D imaging and Z-stack reconstruction of the cell-encapsulated hydrogels.

### 2.15 Statistical analysis

All quantitative data are presented as mean ± standard deviation, with inferential statistics (p-values) used for further analysis. One-way ANOVA was conducted to assess statistical significance, with thresholds set at p < 0.05, p < 0.01, and p < 0.001. Tukey’s HSD test was employed for post-comparisons to determine significant differences between groups. All statistical analyses were performed using GraphPad Prism (version 12.2.2).

## 3. Results and Discussion

### 3.1 TMA synthesis and methacrylation evaluation

TMA was then synthesized through the chemical modification of Taur’s primary amine groups and their replacement with methacryloyl functional groups. The two-step synthesis process has been detailed in the sequence of steps as shown in **Figure 1A**, and the substitution reaction at the molecular scale has been highlighted in **Figure 1B**. An initial substitution reaction at high temperature was performed for 24 h, after which precipitation with 100% ethanol was carried out. The non-reactive GMA was removed by washing and resuspending the reaction mixture three times in ethanol. Ethanol was identified as the bridging solvent due to the miscibility of water and GMA in ethanol. Upon the addition of the TMA reaction solution into ethanol, all liquid components formed one single homogeneous phase. Simultaneously, ethanol served as an anti-solvent for Taur and TMA [46,47]. This enabled the precipitation of TMA due to its different solubility in water and ethanol. After the TMA precipitated in ethanol, centrifugation was then used to extract the precipitated TMA from ethanol, and the supernatant was removed. The precipitate was then resuspended in fresh ethanol to ensure complete removal of any remaining GMA, vortexed and centrifuged. This was repeated three times referred as washing by ethanol three times in this study. Ultimately, the TMA powder was collected, then suspended in a small amount of ethanol and heated overnight at 50°C, so the ethanol was evaporated and thus, reaching the final dried powder.

**Figure 1.**
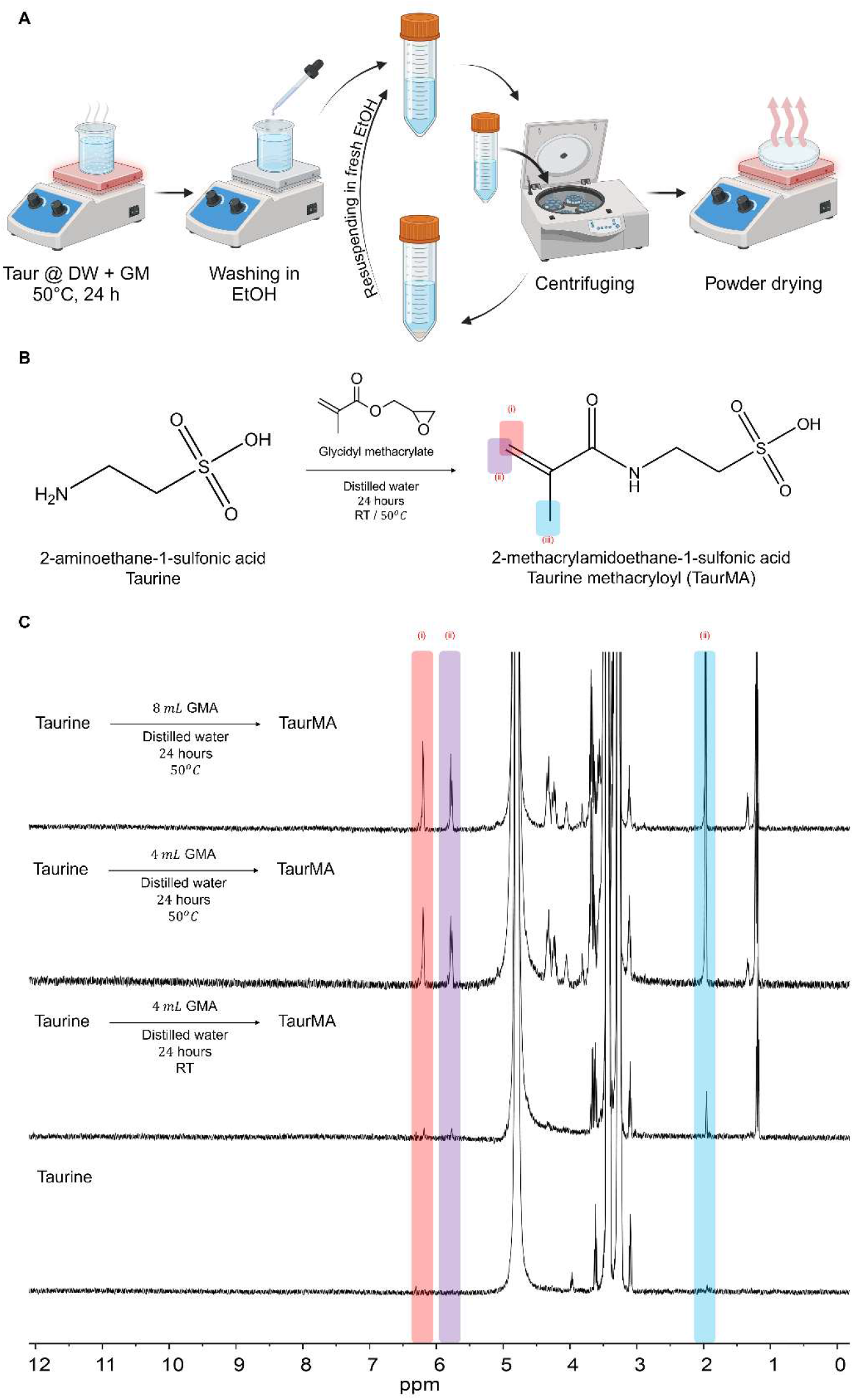
Taur chemical modification. A) Schematic representation of TMA synthesis from Taur (Created in BioRender. Kim, K. (2025) https://BioRender.com/w24y725). B) NMR spectroscopy confirms the substitution of methacrylate functional groups on Taur under different reaction conditions to produce TMA.

Using distilled water as a solvent for the methacrylation reaction; Taur is highly soluble in water. In this step of the reaction, GMA was used as the methacrylation precursor using the previously established optimized protocol for GelMA synthesis by our group [18]. As GMA is not miscible with distilled water; therefore, the solution was kept under constant stirring throughout the reaction at 750 rpm to employ small droplets of GMA in the aqueous phase by emulsion. This was to provide adequate contact surface area of the dissolved Taur molecules to GMA, thus enabling substitution reactions.

To verify methacry late substitution and explore the optimization of reaction conditions, the effects of reaction temperature and GMA concentration on the degree of substitution (DS) were characterized qualitatively and quantitatively via proton nuclear magnetic resonance (^1^HNMR). ^1^HNMR spectra of Taur and TMA synthetized under the different reaction conditions are shown for comparison in **Figure 1C**. With the optimum GelMA synthesis conditions in mind, we examine two reaction temperatures, room temperature and 50°C. Also, two molar ratios of Taur to GMA (1:1.75 and 1:3.5) were used in the methacrylation reaction, corresponding to the ratio utilized for GelMA synthesis.

The methacrylate vinyl-related signals (6.0–6.2, 5.6–5.8 ppm) and signals of methyl groups (1.2–1.8 ppm) were also detected, indicating that methacrylate was successfully substituted on Taur to obtain TMA [18,24,44]. The intensity of methacryloyl chemical group peaks recorded at 1.93, 5.78 and 6.22 ppm as displayed in the NMR data was higher at 50°C reaction condition relative to room temperature. In addition, raising the Taur:GMA molar ratio from 1:1.75 to 1:3.5 substantially improved the DS, with an increase in substitution efficiency, which suggests that more amine groups were modified with methacryloyl functional groups. As a result, 50°C and a 1:3.5 molar ratio was selected as the optimal conditions to perform this reaction.

### 3.2 Physiomechanical properties of the hydrogels

Due to several chemical reasons, regardless of functionalizing Taur and synthesizing TMA, it may not form a stable hydrogel network. Unlike traditional hydrogel precursors, such as GelMA or PEGDA, which contain multiple reactive sites, TMA only has one methacrylate group per molecule, meaning it cannot form a crosslinked network on its own. Additionally, Taur contains a sulfonic acid (-SO_3_H) group, which is highly hydrophilic. This strong hydrophilicity can prevent chain entanglement and network stabilization, leading to dissolution rather than gelation. Instead, it undergoes linear polymerization, resulting in water-soluble or weakly crosslinked structures that dissolve or fail to form a stable gel. Therefore, a polymeric network presence is necessary to make a stable network that contains TMA in it. Among several photocrosslinkable hydrogels, GelMA is a versatile photocrosslinkable hydrogel ideal for biomedical and tissue engineering due to its biocompatibility, and tunable mechanics. It retains RGD peptide sequences for cell adhesion and allows precise crosslinking under light exposure, making it excellent for 3D cell culture, tissue regeneration, and bioprinting [48,49]. Hence, within the designed system, GelMA served as base hydrogel, and TMA was co-polymerized with GelMA to get the final hybrid bioactive hydrogel network **(Figure 2A)**. After fabricating the main structure of the hybrid hydrogel, the physical properties of the hydrogels containing 5% GelMA with different concentrations of Taur and TMA were evaluated. **Figure 2A** represents the mechanical properties of the hybrid hydrogels containing Taur and TMA. It’s shown that by adding the Taur and TMA, the mechanical properties of 5% GelMA decreased from 11.4 ± 2.7 kPa to 6 kPa and 7 kPa for Taur and TMA groups, respectively. In GelMA photocrosslinking, the process relies on the generation of free radicals (using LAP as a photoinitiator) to initiate methacrylate group polymerization. Taur contains a sulfonic acid (-SO_3_H) group, which has been reported to act as a free radical scavenger by neutralizing reactive oxygen species (ROS) and polymerization radicals [50–54]. The presence of Taur and TMA scavenges these radicals, thereby reducing the number of available free radicals for crosslinking which limits the extent of crosslinking, resulting in a weaker and less dense polymer network. As shown in **Figure 2B(i)**, a further increase in Taur concentration did not result in a significant change in the compression modulus **(Table S1)**. This consistency in mechanical properties may be attributed to the accumulation of Taur crystals within the GelMA walls of the hydrogel microstructure, as illustrated in **Figure 4A**, leading to stiffer microstructure. However, **Figure 2B(ii)** supports the observation that increasing TMA concentration led to a slight increase in the compression modulus, from 5.4 ± 0.93 kPa at 1% TMA to 7.4 ± 1.1 kPa at 10% TMA. This increase is likely due to a higher degree of covalent crosslinking, resulting from the greater availability of methacry late groups in the hydrogel network.

**Figure 2.**
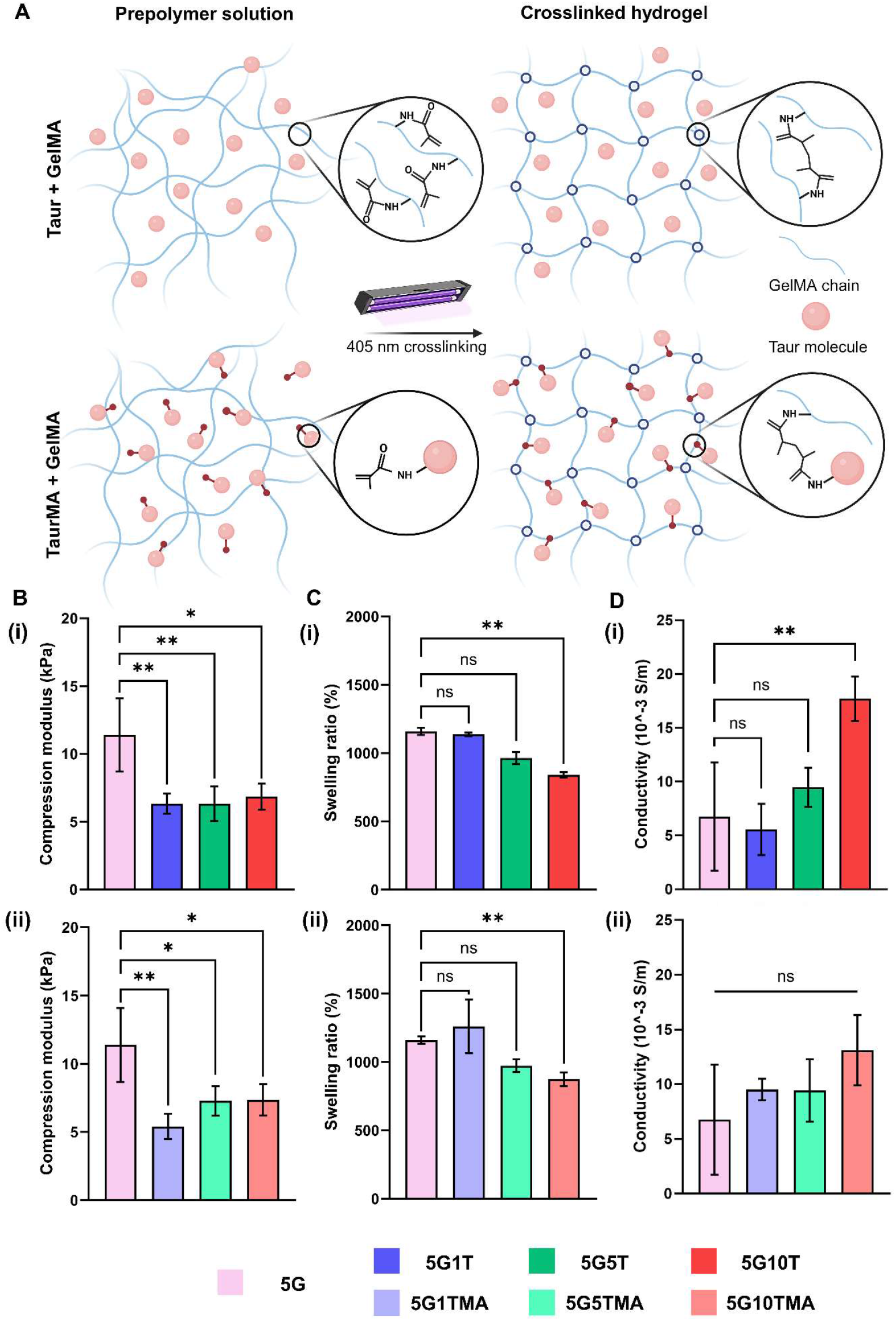
Mechanical and physical characterization of bioinks containing Taur and TMA. A) Schematic illustration of the chemical interaction between Taur, TMA, and GelMA before and after photocrosslinking (Created in BioRender. Kim, K. (2025) https://BioRender.com/o43t997). B) Mechanical properties of 5% GelMA containing 0, 1, 5, and 10% (i) Taur and (ii) TMA. C) Swelling ratio of 5% GelMA containing 0, 1, 5, and 10% (i) Taur and (ii) TMA. DJ Conductivity of 5% GelMA containing 0, 1, 5, and 10% (i) Taur and (ii) TMA. For all experiments, n=5.

**Figure 3.**
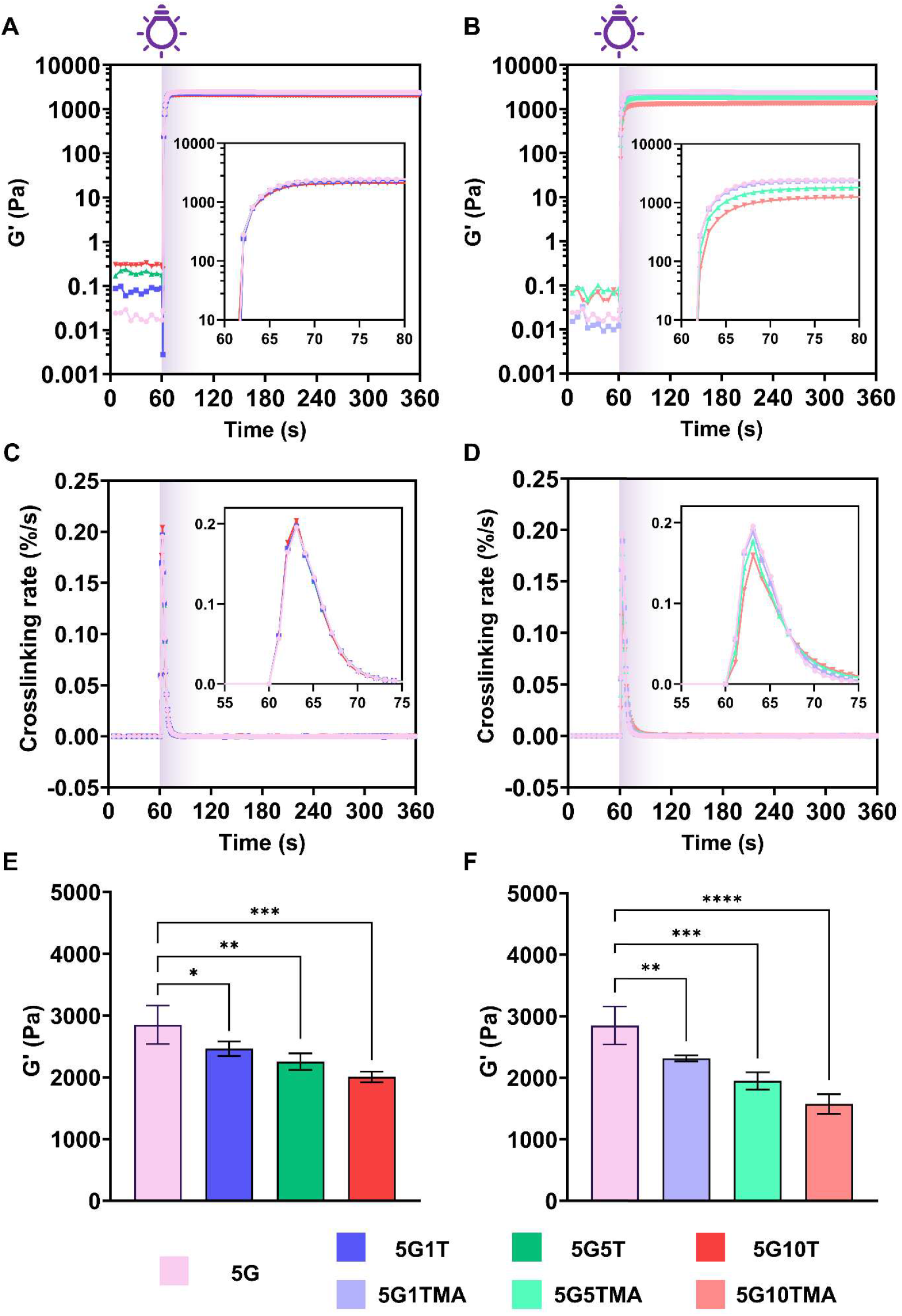
Photo-crosslinking kinetics of Taur- and TMA-containing GelMA hydrogels. A, B) Photo-crosslinking kinetics of GelMA hydrogels containing (A) Taur and (B) TMA. C, D) Variation in crosslinking rate for GelMA hydrogels with (C) Taur and (D) TMA. E, F) Saturated storage modulus of crosslinked GelMA hydrogels with (E) Taur and (F) TMA. For all experiments, n = 3.

**Figure 4.**
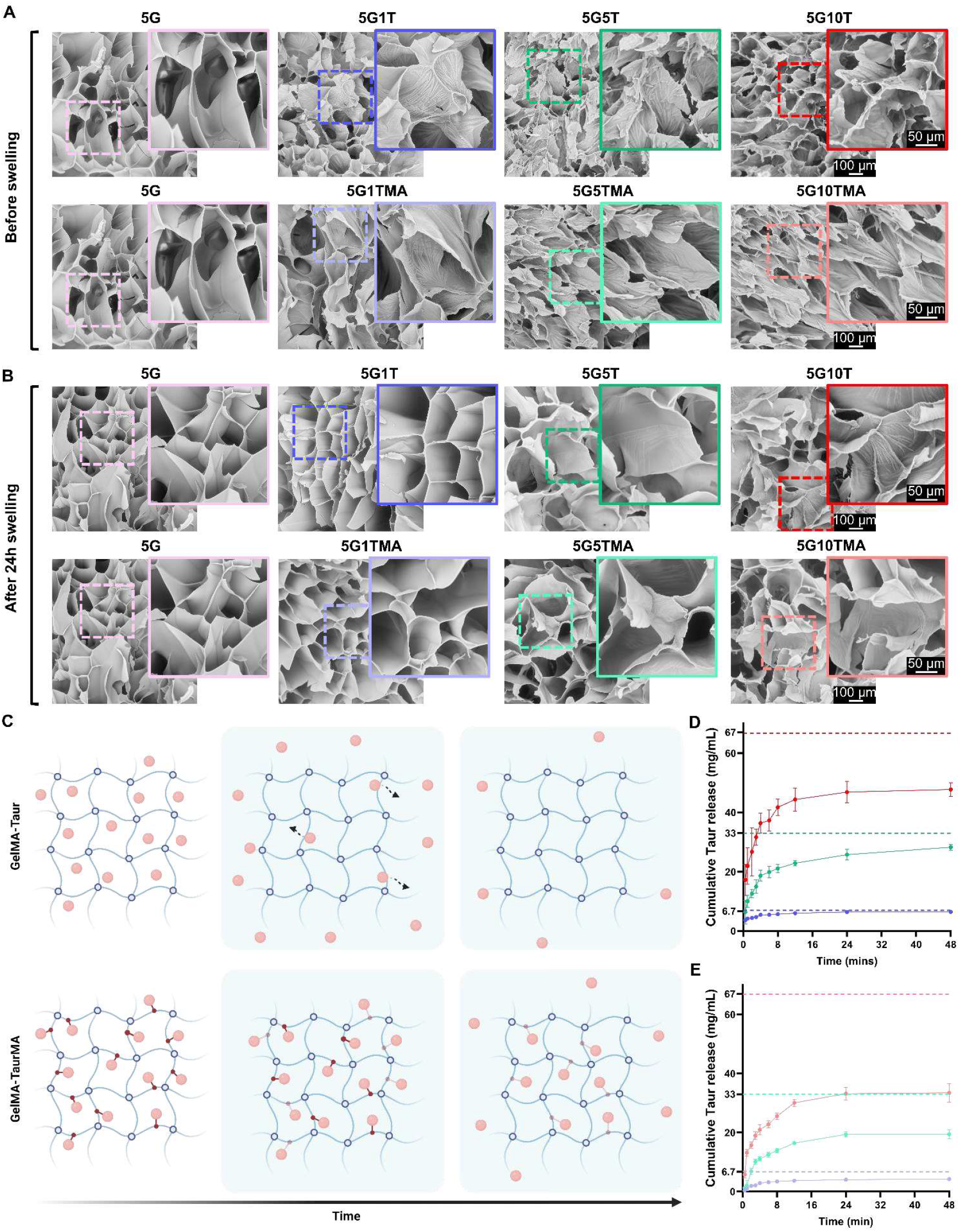
Microstructural analysis and release kinetics of Taur- and TMA-containing GelMA hydrogels. A) SEM images showing the microstructure ofGelMA hydrogels with different concentrations of Taur and TMA after photo-crosslinking. B) SEM images showing the microstructure of GelMA hydrogels with different concentrations of Taur and TMA after 24 h of swelling. C) Schematic illustration of the release mechanism of GelMA hydrogels containing Taur and TMA over time in PBS (Created in BioRender. Kim, K (2025) https://BioRender.com/x60b534). D) Release kinetics of GelMA hydrogels containing Taur over time. The horizontal lines indicate the initial Taur concentration in the samples. E) Release kinetics of GelMA hydrogels containing TMA over time. The horizontal lines indicate the initial TMA concentration in the samples. For all experiments, n=5.

In addition to mechanical properties, swelling characteristics play a crucial role in solute diffusion and nutrient circulation within the scaffold, affecting the transport of essential molecules such as oxygen, nutrients, and growth factors, which are critical for cell survival and tissue regeneration [55]. **Figure 2C** illustrates a decreasing trend in the swelling ratio for both Taur and TMA experimental groups. Hydrogels exhibit complex swelling behavior influenced by ionic interactions and network structure. It is well established that an increase in the ionic strength in the hydrogel environment leads to a reduction in swelling capacity [56–58]. The charged sulfonic acid group (-SO_3_H) in Taur can enhance ionic interactions within the hydrogel network, leading to a more compact structure. This structural compaction, confirmed through microstructural evaluation in **Figure 4**, results in a decreased swelling ratio—from 1160.02 ± 26.52% in the 5G sample to 841.61 ± 19.45% in 5G10T and 874.59 ± 49.52% in 5G10TMA.

However, at lower Taur and TMA concentrations, the swelling ratio does not differ significantly from the control sample **(Figure 2C(i)** and **(ii))**. This aligns with the structural compactness observed in **Figure 4**, where the water uptake capacity remains similar. This behavior may be attributed to the high hydrophilicity of Taur, which counterbalances the network compaction by facilitating increased water absorption.

**Figure 2D (i)** shows the electrical conductivity measurements of GelMA hydrogels modified with Taur and TMA. The electrical conductivity measurements were carried out as per the protocol outlined in section 2.4. Electrical conductivity is a critical property for supporting excitable cell types, such as neurons and muscle cells, as it facilitates the transmission of electrical signals essential for their function and maturation. Electroconductive hydrogels emulate the native ECM by providing electrical cues that regulate cellular behavior and promote proper development and physiological activity [59,60]. The pristine GelMA (5G) exhibits the lowest conductivity (∼6×10^−3^ S/m), while after the incorporation of Taur, the highest electrical conductivity was observed in the case of 5G10T (∼17 × 10^−3^ S/m). In contrast, samples 5G1T and 5G5T showed nonsignificant differences in their electrical conductivity values when compared to the control sample 5G. Next, the incorporation of TMA also leads to an increase in the electrical conductivity of the samples **(Figure 2D (ii))**. Specifically, the 5G10TMA shows enhancement in electrical conductivity (∼13×10^−3^ S/m), highlighting the effectiveness of covalently grafted sulfonic acid groups in facilitating ionic transport. Although 5G10TMA demonstrates the highest average conductivity, no statistically significant differences are observed among 5G1TMA, 5G5TMA, and 5G10TMA groups. This suggests a plateau effect, likely due to ionic crowding or saturation of charge transport pathways at higher TMA concentrations. The difference in electrical conductivity between Taur and TMA is obvious as Taur is not chemically integrated into the polymer network; the ions have higher mobility and, therefore, result in higher conductivity values compared to well-attached TMA **(Figure S1)**. Similar behaviour was observed by Wu et al. [61], wherein they observed that charge transport is more efficient in pure GelMA as compared to GelMA-Pani crosslinked sample. This is likely due to unrestricted diffusion of ions, which becomes hindered upon crosslinking with Pani. Therefore, this helps in understanding the difference between the electrical conductivity values of the Taur and TMA groups. Moreover, optical transparency is another hydrogel’s essential property, particularly in biomedical applications such as cell encapsulation, tissue engineering, and optical biosensing, where real-time visualization of internal structures or cellular activity is essential. Moreover, numerous studies have highlighted the impact ofhydrogel transparency on the resolution and precision oflight-based bioprinting techniques [62–64]. The transparency of the hydrogels was evaluated by using UV-Vis method. All samples showed high transparency in the range of visible light; there was no significant difference between the two groups **(Figure S*2*A** and **B)**.

**Figure 5.**
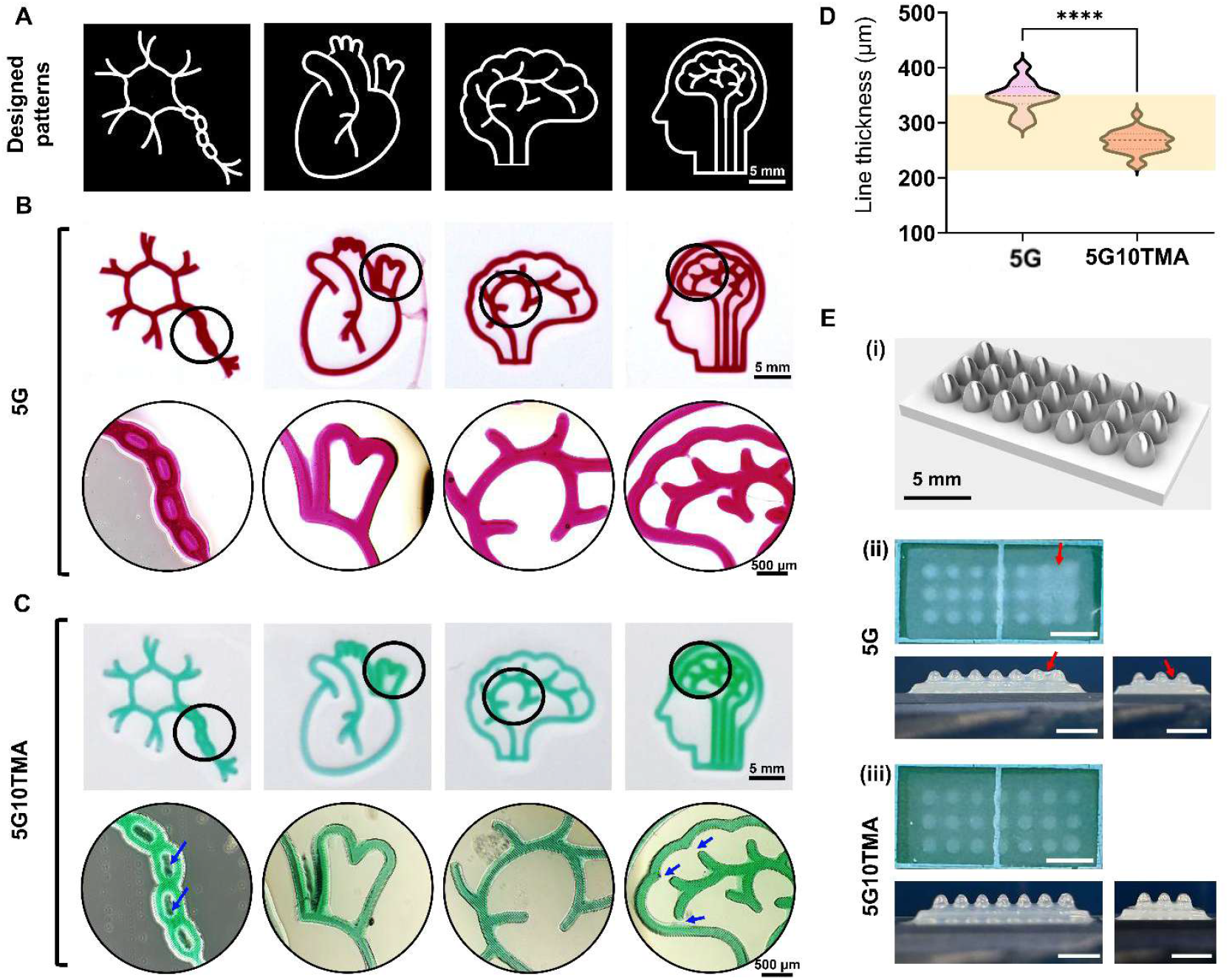
Printing evaluation of TMA containing GelMA hydrogels. A) Patterns used for photopatterning of bioinks. B) Photopatterned and stained structures after crosslinking 5G hydrogel (crosslinking time = 10 s) and images captured using 4 × brightfield imaging. C) Photopatterned and stained structures after crosslinking 5G10TMA hydrogel (crosslinking time= 10 s) and images captured using 4 × brightfield imaging. D) Line thickness measurements of the different photopatterned structures, with the light beam thickness range used for photo-crosslinking highlighted in yellow. E) Milipillar arrays 3D printing, (i) CAD model, (ii) 3D printed structure using 5G bioink, (iii) 3D printed structure using 5G10TMA (scale= 5mm).

### 3.3 Photocrosslinking characteristics

**Figure 3A** illustrates the evolution of the storage modulus (G’) over time during the photo-crosslinking process. The stiffness of the hydrogel as crosslinking progresses is depicted by G’. All the samples exhibited a plateau in the measured G’ curves at the end of the illumination period, implying saturation in the degree of photo-crosslink:ing. Pristine GelMA (5G) demonstrated the highest final G’, indicating the strongest hydrogel network formation. In comparison, 5G1T exhibited a slight reduction in G’, while 5G5T and 5G10T showed progressively lower values, with 5G10T having the lowest stiffness. These observations suggest that Taur, at higher concentrations, interferes with the GelMA photo-crosslink:ing process. The hydrophilic nature of Taur likely disrupts the GelMA network by diluting the polymer matrix, leading to decreased mechanical stiffness of the resulting GelMA hydrogels [65], along with the free radical scavenging nature of Taur as described earlier. The inset of the graph highlights the rapid increase in G’ during the early stages of photo-crosslinking, further emphasizing the effect of Taur on initial hydrogel network formation. The change in the photo-crosslinking kinetics of GelMA bioinks modified with varying concentrations of TMA was also evaluated under similar photo-crosslinking conditions. **Figure 3B** illustrates the evolution of the storage modulus over time, which represents the stiffness of the hydrogel during photo-crosslinking. 5G sample exhibited the highest G’, indicating the strongest polymeric network formation, while hydrogels with increasing TMA concentrations (5G1TMA, 5G5TMA, and 5G10TMA) demonstrated progressively lower G’. This reduction in stiffness at higher TMA concentrations is attributed to the disruption of polymer entanglements and the introduction of hydrophilic methacrylate-functionalized Taur groups, which interfere with the crosslinking process. Further, the inset highlights the rapid rise in G’ during the early phase (55–80 seconds), with a slight increase observed for increasing TMA concentrations.

**Figure 3C** depicts the crosslinking rate as a function of time during the gelation process. The peak crosslinking rate represents the speed of network formation. 5G control sample exhibited the highest peak crosslinking rate, indicating rapid gelation. In comparison, 5G1T showed a marginally reduced rate, while 5G5T and 5G10T demonstrated significantly slower rates, with 5G10T having the lowest. The decrease in crosslinking rate with increasing Taur concentration is consistent with Taur’s interference in the photopolymerization process. Taur introduces hydrophilic groups, increasing water retention in the system and reducing crosslinking efficiency. The inset highlights the peak crosslinking rates within the critical time window, showing the diminished crosslinking dynamics at higher Taur concentrations. Similarly, **Figure 3D** presents the crosslinking rate (%/s) as a function of time, capturing the dynamics of network formation for GelMA-TMA hydrogels. GelMA without TMA (5G) exhibited the fastest crosslinking rate, while the rates decreased with increasing TMA concentrations. The slower rates for 5G5TMA and 5G10TMA suggest that higher TMA concentrations introduce steric hindrance and reduce crosslinking efficiency, correlating with the lower stiffness in the resulting GelMA-TMA composite hydrogels. Further, it should be noted that this effect of taur/TMA in GelMA is due to the scavenging effect [66,67], wherein higher concentrations may overly quench the photoinitiated radicals, thereby reducing the crosslinking efficiency. The excess TMA can also contribute to increased hydrophilicity, leading to higher water uptake, matrix swelling, and plasticization, which soften the hydrogel structure. Additionally, the inset highlights the peak crosslinking rates within the critical gelation window (55–75 seconds), further supporting the conclusion that TMA disrupts crosslinking dynamics.

**Figure 3E** compares all hydrogel formulations’ final storage modulus (G’) after crosslinking. 5G sample had the highest G’, serving as the benchmark for stiffness. The addition of 1% Taur (5G1T) resulted in a slight but statistically significant reduction in G’ *(p* < *0*.*05)*, while the hydrogel with 5% Taur (5G5T) showed a more pronounced decrease *(*p* < *0*.*01)*. 5G10T sample exhibited the lowest G’, with a highly significant difference from 5G *(**p* < *0*.*001)*. These results indicate that increasing Taur concentrations progressively weaken the hydrogel network due to interference with the photocrosslinking process, which is aligned with the mechanical properties analysis mentioned previously. This trend highlights the tunability of GelMA hydrogels, where stiffness can be modulated based on Taur content. Subsequently, **Figure 3F** compares the final storage modulus (G’) of the crosslinked GelMA-TMAhydrogels. Similarly, 5G showed the highest G’, serving as the baseline, while 5G1TMA, 5G5TMA, and 5G10TMA exhibited progressively lower final values of G’. The statistical significance of these reductions (p < 0.01 for 5G1TMA, *p* < *0*.*001* for 5G5TMA, and p < 0.0001 for 5G10TMA) highlights the impact of TMA concentration on crosslinked hydrogel stiffness. The decrease in G’ can be attributed to the disruption of GelMA network caused by TMA incorporation. This trend indicates that both Taur/TMA-modified GelMA hydrogels offer tunable mechanical properties, with higher Taur/TMA concentrations leading to hydrogels suitable for applications requiring flexibility and biocompatibility, such as engineered soft tissues. The viscoelastic behavior of photocrosslinked hydrogel formulations, including GelMA, taurine-incorporated GelMA, and TMA with GelMA, was analyzed across a range of oscillatory frequencies (see **Figure S3)**. In all compositions, the storage modulus (G’) remained higher than the loss modulus (G”) throughout the frequency range, indicating dominant elastic (solid-like) behavior. This further confirms the complete crosslinking of the hydrogel samples. As frequency increased, all samples demonstrated a moderate rise in G’ and G”, consistent with viscoelastic hydrogels where polymer chain motion becomes restricted under rapid deformation.

### 3.4 Release evaluation

To investigate the influence of Taur and TMA on the microstructure of GelMA, different combinations were evaluated using SEM. **Figure 4A** shows the microstructure of pristine GelMA alongside samples containing Taur and TMA. In both groups, it is evident that introducing Taur and TMA into GelMA leads to a decrease in pore size. This decreasing trend is more pronounced in samples containing 5% or higher concentrations of Taur or TMA. This observation aligns with the swelling analysis, which also showed that increasing Taur and TMA concentration led to a decrease in swelling ratio. The reduction in pore size could be attributed to several factors. Both Taur and TMA contain sulfonic acid (-SO_3_H) and amine (-NH2) groups, which can form hydrogen bonds and electrostatic interactions with the GelMA matrix. These interactions can tighten the hydrogel network even in the absence of covalent crosslinking, contributing to pore size reduction. In the case of TMA, due to the presence of methacry late groups, the crosslinking density increases, resulting in a more significant decrease in pore size. Furthermore, the charged functional groups in Taur and TMA may induce electrostatic attractions between polymer chains, effectively drawing them closer together, enhancing molecular packing and limiting the formation of large pores.

After 24 hours of swelling in PBS, the microstructure of the samples was examined again using SEM, as shown in **Figure 4B**. It is evident that, except for the pristine GelMA, the Taur- and TMA-containing GelMA samples exhibited increased pore sizes after extended swelling. Over time, hydrogels can undergo network relaxation or chain rearrangement, and the influx of water during swelling can push polymer chains apart. Additionally, some amount of Taur and TMA may be released into the PBS during this process. In the absence of these molecules, the network faces less internal resistance to expansion, allowing for greater chain mobility and pore enlargement.

Beyond their influence on pore size and structural compactness, Taur and TMA leave visible signs of their presence in the GelMA structure, as shown in **Figure 4A** and **B**. The addition of Taur and TMA results in reinforcement of the pore walls with Taur- and TMA-derived fibrils. These lamellar precipitates are more prevalent in Taur-containing hydrogels, likely due to the lack of covalent bonding between Taur and GelMA. During crosslinking, phase separation may occur, dragging Taur molecules into regions being crosslinked last, which then crystallize on the hydrogel walls. In contrast, TMA provides a more homogeneous structure due to its ability to form covalent bonds with GelMA, embedding it within the GelMA walls and leading to fewer visible crystalline structures.

SEM elemental mapping confirms the distribution of Taur and TMA within the GelMA structure, as illustrated in **Figure S4A** and **Figure S5A**, respectively. **Figure 4B** suggests that Taur and TMA crystals on the GelMA surface become less apparent after swelling, likely due to their release into the medium. However, the amount of Taur or TMA remaining cannot be solely determined from visible crystalline residues in SEM images, as some may still be blended within the GelMA walls, requiring EDS analysis for confirmation.

After 24 hours of swelling, a portion of Taur and TMA was released into the surrounding PBS. As shown in **Figure 4B**, in the 5G1T and 5G1TMA samples, no visible Taur or TMA crystals remain, indicating that most of the additive was washed out. EDS quantitative elemental analysis, presented in **Table S2** and **Table S3**, supports this. For the 5G1T sample, sulfur (S) content decreased from 2.43% to 0%, suggesting that Taur was almost entirely depleted from the scaffold and was no longer detectable by EDS. In contrast, in the 5G1TMA sample, 0.28% sulfur remained after swelling, indicating that some TMA is still retained, likely trapped within the GelMA walls, as no surface crystals were visible. Similarly, for samples with greater than 1% Taur or TMA, EDS analysis reveals greater retention of sulfur in the TMA group, due to covalent bonding between TMA and the GelMA backbone, which limits TMA release into the medium **(Table S2** and **Table S3)**. SEM elemental mapping images post-swelling **(Figure S4B** and **Figure S5B)** also confirm that sulfur content is higher in TMA-containing samples after 24 hours of swelling.

As discussed, both Taur and TMA are partially released during swelling. The suggested release mechanism is illustrated in **Figure 4C**. In the GelMA-Taur system, Taur molecules are physically entrapped and stabilized by weak hydrogen bonding and electrostatic interactions. Due to Taur’s high solubility in water, it dissolves in PBS and leaves the hydrogel matrix easily. In contrast, in the GelMA-TMA system, covalent bonds between TMA and GelMA branches hinder release. Initially, surface-bound TMA crystals dissolve in PBS, but the covalently bonded TMA takes longer to be released. Over time, as some covalent bonds break, loosely bound TMA molecules may gradually leave the structure, while the majority remains embedded within the GelMA walls.

To investigate the release kinetics of Taur and TMA over an extended swelling period, a release experiment was conducted, and Taurine-ELISA was used to quantify the release of both compounds. The results are presented in **Figure *4*D** and **E**. In Tam-containing samples, a noticeable initial burst release was observed, indicating that a large portion of Taur crystals, previously observed on the GelMA wall surface in SEM images **(Figure *4*A)**, were loosely attached and readily released into the medium **(Figure *4*D)**. This was followed by a continuous release at a higher rate compared to the TMA-containing samples, as shown in **Figure *4*D** and **E. Figure *4*E**, supported by EDS analysis **(Table S*2)***, shows that up to 96% of the initially encapsulated Taur was released after 48 hours of PBS immersion in the 5G1T sample. In contrast, in the 5G1TMA sample, approximately 40% of TMA remained after 48 hours. Additionally, **Figure *4*E** demonstrates the sustained release profile of TMA, with slower release rates across all TMA-containing groups compared to Taur-containing ones. Moreover, TMA and Taur retention levels were over 51% in 5G10TMA and 28% in 5G10T, respectively, after 48 hours, indicating that the presence of methacrylate groups and covalent bonding between TMA and the GelMA backbone contributed to controlled release and long-term retention of TMA within the hydrogel structure.

### 3.5 Photo patterning and 3D printing evaluation

As mentioned previously, Taur can act as a free radical scavenger when added to the GelMA prepolymer. Its influence on mechanical properties has been investigated, and its scavenging effect has been confirmed.

Therefore, it is important to evaluate whether the presence ofTMA affects printability. To assess this, 5G and 5G10TMA (GelMA with the highest TMA concentration) were selected for photo-patterning and 3D printing. **Figure 5** presents the various designs used for photo-patterning GelMA hydrogels, while **Figure 5B** and **C** display the crosslinked patterns alongside microscopic images highlighting the details of the printed structures. In both groups (5G and 5G10TMA), different structures were successfully crosslinked with high shape fidelity after washing away the uncrosslinked prepolyrner solution. However, brightfield images reveal some differences between 5G and 5G10TMA. As shown with orange arrows in **Figure 5B** and **C**, the hollow sections in the chain structure were over-crosslinked in 5G, while in the presence of TMA, these hollow sections were successfully retained. Since LAP was used as the photo-initiator, it undergoes photo-cleavage upon exposure to 405 nm light, generating free radicals that initiate GelMA crosslinking via rnethacry late groups. Meanwhile, TMA contains a highly reactive sulfonic acid (-SO_3_H) group, which can interact with free radicals by donating electrons [68], effectively neutralizing them before they initiate crosslinking. This reaction competes with GelMA polymerization, potentially reducing the number of successful crosslinks, leading to lower crosslink density and a softer hydrogel. Previous sections have shown that even in the presence of Taur or TMA, stable structures are still formed over time, confirming that a sufficient degree of crosslinking is maintained. This suggests that TMA regulates the crosslinking density, preventing over-crosslinking and resulting in sharper, more defined structures. This effect can also be seen in other structures, such as the fourth one, where blue arrows highlight distinct uncrosslinked areas between two branches, preventing undesired crosslinking.

Additionally, the crosslinked 5G10TMA hydrogels **(Figure 5C)** appear dimmer compared to the photo-patterned 5G constructs **(Figure 5B)** after staining. This could be due to the lower swelling ratio of TMA-containing groups **(Figure 2C(ii))**, which may reduce dye absorption.

Furthermore, the thickness of the patterned structures was evaluated **(Figure 5D)**. The yellow highlighted region in the graph represents the acceptable light beam thickness range, demonstrating that 5G10TMA crosslinked structures fall within this range. In contrast, 5G samples produced significantly thicker lines, likely due to uncontrolled crosslinking kinetics and over-crosslinking. To further investigate the potential of the developed bioink for DLP 3D printing, both 5G and 5G10TMA were processed using a DLP 3D printer. **Figure 5E(i)** presents the CAD model containing micropillar structures, which was used to assess hydrogel printability. **Figure 5E(ii)** and **(iii)** show the 3D-printed structures fabricated from 5G and 5G10TMA, respectively. Similar to the photo-patterning experiments, over-crosslinking was observed in the 5G crosslinked hydrogel, as indicated by the red arrows **(Figure 5E(ii)**). In contrast, 5G10TMA demonstrated more controlled crosslinking, resulting in sharper, more defined micropillars during the DLP 3D printing process **(Figure 5E(iii))**.

Moreover, the 3D constructs printed with 5G10TMA appeared more transparent, whereas 5G hydrogels exhibited a cloudy appearance after crosslinking. This could be attributed to the hydrophilic sulfonic acid (-SO_3_H) groups in TMA, which may enhance polymer dispersion, reduce phase separation during crosslinking, and lead to a more homogeneous polymer network, ultimately reducing light scattering.

To further evaluate the potential of the TMA-containing bioink for 3D printing, twisted pyramid structures **(Figure S6B)** were printed using both 5G and 5G10TMA inks. The brightfield images in **Figure S6C** confirm the successful fabrication of these structures with both bioinks. Additionally, **Video S1** further supports the successful 3D construction of twisted pyramids using GelMA-based hydrogels. Additionally, to assess the resolution of DLP 3D printing, the CAD model shown in **Figure S6A** was used to print various structures using 5G and 5G10TMA, followed by thickness measurements of each printed line to compare printing resolution. The results, illustrated in **Figure S7A** and **B**, show that for all lines in the given design, the actual printed structures produced thinner lines when using 5G10TMA compared to 5G under the same crosslinking conditions, highlighting the potential for achieving higher 3D printing resolution.

### 3.6 Biocompatibility characteristics

For the biological study, C2C12 myoblasts were encapsulated within the 3D GelMA hydrogel matrix, both with and without Taur and TMA. To assess the cytotoxicity of TMA, a live/dead assay was performed over a one-week culture period on the control hydrogel and hydrogels containing the highest concentrations of Taur and TMA. **Figure 6A** presents live cells (green) and dead cells (red) at two different magnifications, allowing both viability assessment and morphological analysis. The predominant green fluorescence across all scaffold groups (5G, 5G10T, and 5G10TMA) indicates high cell viability, which can be attributed to sufficient nutrient transport through the porous scaffold and the non-toxic nature of Taur and TMA. Additionally, quantitative viability analysis **(Figure 6B)** showed no significant difference in cell viability across the different scaffolds over seven days of culture.

**Figure 6.**
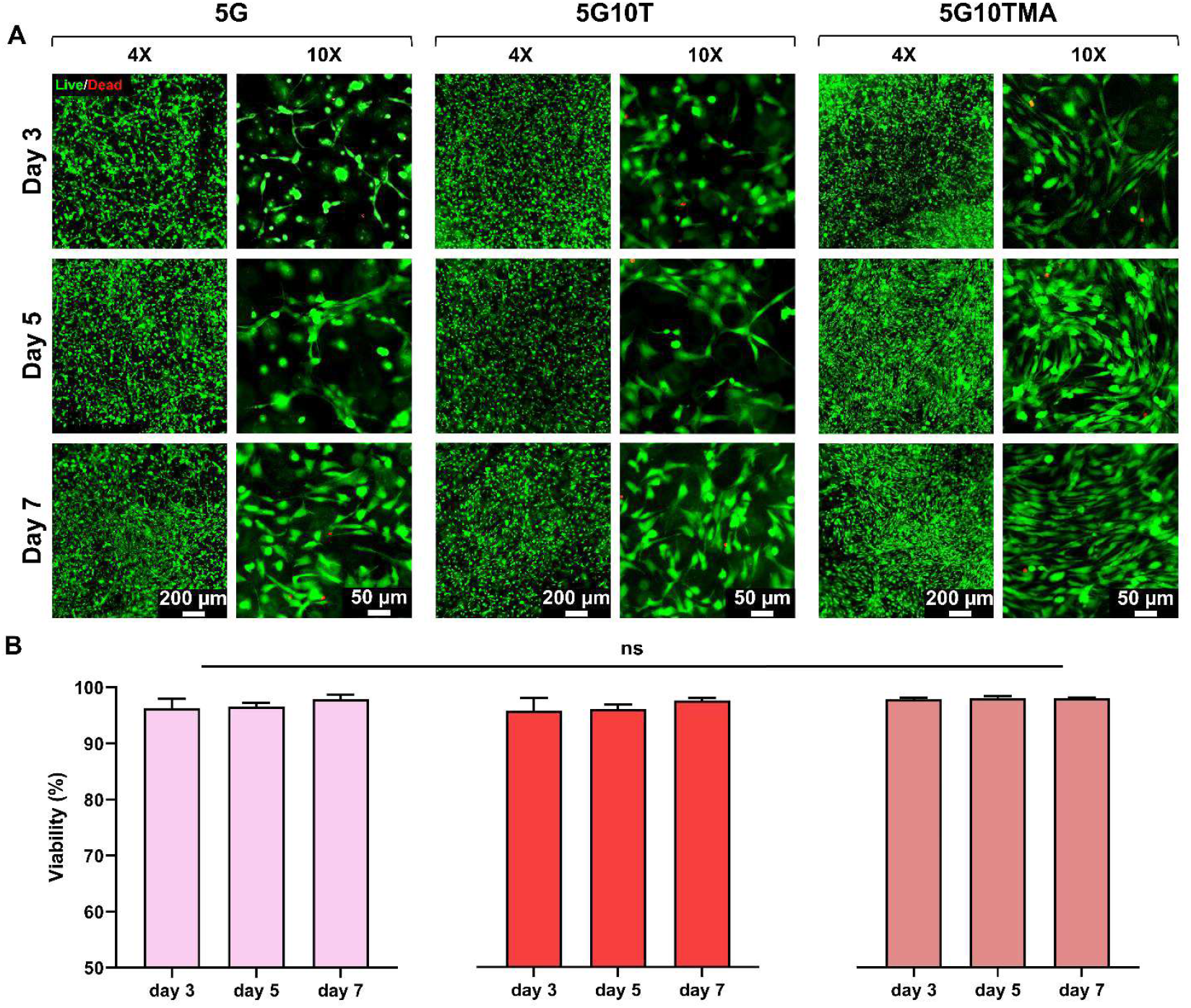
Viability evaluation of 3D encapsulated C2C12 myoblasts in 5G, 5G10T, and 5G10TMA hydrogels. A) Flourescence images of live/dead stained samples cultured for 3, 5, and 7 days. B) Quantitative analysis of cell viability in different hydrogel samples over 7 days of culture.

Beyond cytocompatibility of Taur and TMA, higher magnification imaging provides insights into cell morphology within the 3D scaffold network. As previously reported, Taur plays a significant role in myoblast proliferation and differentiation via the PI3K/Akt signaling pathway [69–71]. A similar effect was anticipated in this study, particularly within the 3D hydrogel structure containing Taur and TMA. **Figure 6A** qualitatively suggests that after seven days of culture, cells proliferated at a higher rate in Taur- and TMA-containing groups compared to the control. However, cells in the TMA group matured more rapidly, which may be due to the chemical integration of TMA into the GelMA hydrogel via covalent crosslinking. This covalent incorporation ensures a more homogeneous polymer network, preventing phase separation or aggregation effects that could occur in the 5G10T group. As a result, TMA provides a more uniform spatial distribution of cell adhesion sites, fostering enhanced cell interactions and proliferation. Additionally, in 5G10TMA, the covalently attached TMA remains embedded in the network longer, exerting a sustained bioactive effect on cells, as demonstrated in the previous section. On the other hand, when comparing 5G10T to the control group, proliferation appeared slightly higher, but myoblasts exhibited a more elongated morphology and increased intercellular connections. This structural organization suggests that cells in 5G10T were transitioning toward myotube formation more rapidly, highlighting the potential role of Taur in accelerating myogenic differentiation.

### 3.7 Cell morphology, myotube formation and image analysis

The process of multinucleated myotube formation begins with the adhesion of mononuclear myoblasts, followed by membrane reorganization **(Figure 7A)**. The rate of myoblast fusion is largely influenced by cell proliferation and migration [72,73]. However, under natural physiological conditions, myoblast fusion can be restricted. In such cases, different external cues such as electrical stimulation, growth factors, and bioactive small molecules can modulate the tissue microenvironment, enhancing cell proliferation and differentiation, ultimately facilitating the fusion of myoblasts into multinucleated myotubes. To assess the effect of Taur on myoblast differentiation and fusion, C2C12 cells were encapsulated in 5% GelMA as the control, along with groups containing varying concentrations of TMA. To examine cell morphology and quantify myotube formation, F-actin staining was performed.

**Figure 7.**
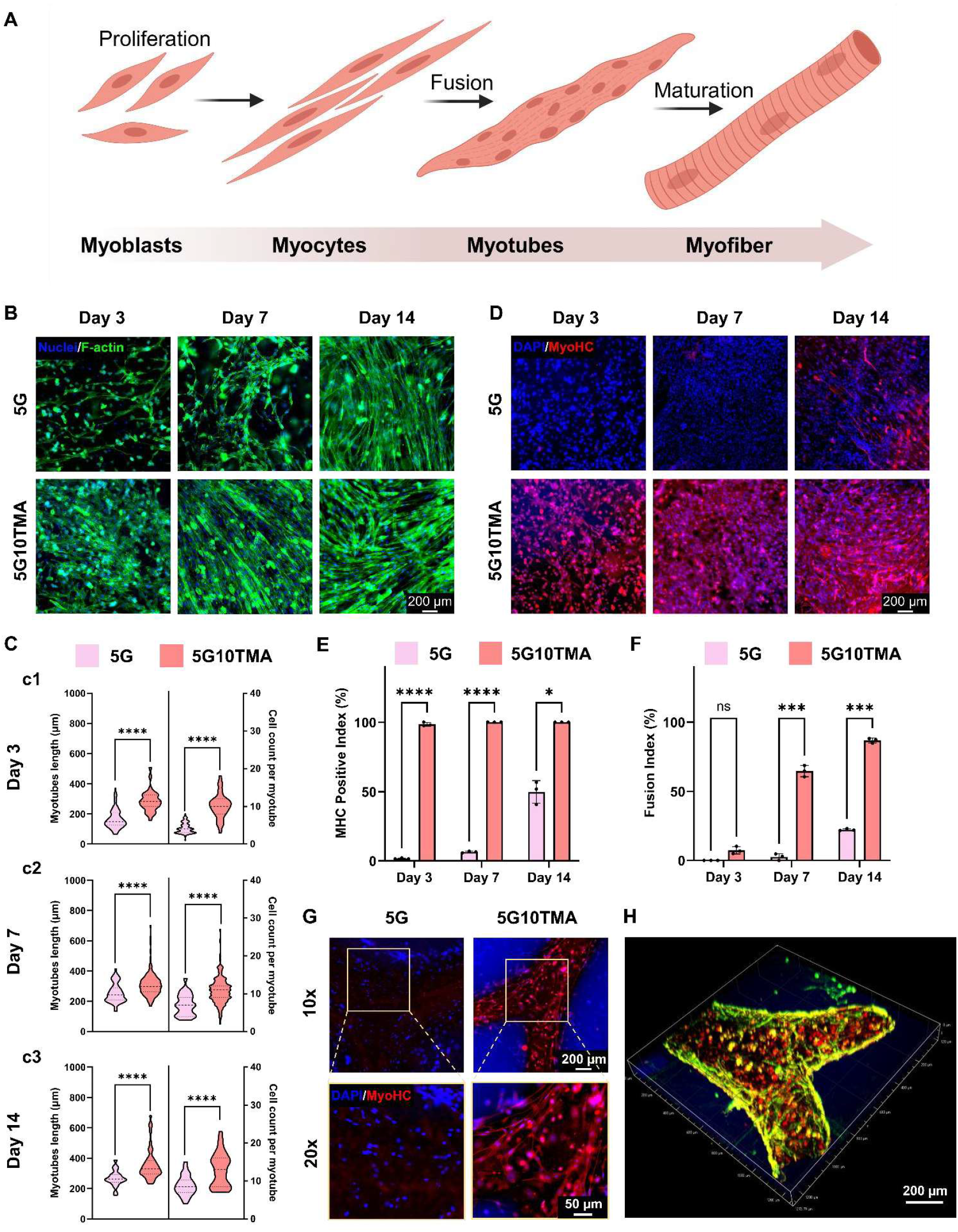
Morphological analysis and myogenic evaluation of 3D-encapsulated C2C12 myoblasts. A) Schematic representation of the differentiation process of myoblasts into myotubes (Created in BioRender. Kim, K. (2025) https://BioRender.com/r56n146). B) F-actin/DAPI immunostaining of 5G and 5G10TMA cell-laden scaffolds for 14 days of culture. C) Image analysis evaluating myotube length and cell count per myotubes for 5G and 5G10TMA cell-laden scaffolds at cl) day 3, c2) day 7, and c3) day 14 of culture. D) MHC/DAPI immunostaining of 5G and 5G10TMA cell-laden scaffolds for 14 days of culture. E) MHC-positive index of 5G and 5G10TMA cell-laden scaffolds over 14 days of culture. F) Fusion index of 5G and 5G10TMA cell-laden scaffolds over 14 days of culture. G) MHCIDAP I immunostaining of 3D bioprinted 5G and 5G10TMA cell-laden scaffolds after 7 days of culture. H) Confocal 3D imaging Phalloidin/MHC/DAPI immunostaining of 3D bioprinted 5G10TMA cell-laden scaffolds after 7 days of culture.

As shown in **Figure 7B**, the proliferation and morphological evaluation of C2C12 myoblasts over 14 days of culture is presented. TMA has been shown to positively influence cell proliferation, as evidenced by the significantly higher Actin-stained area in the 5G10TMA sample as early as day 3. Previous studies have demonstrated that Taur stimulates protein synthesis and myoblast proliferation through the PI3K-ARID4B-mTOR pathway [74–76]. After 7 days of culture, C2C12 cells in the TMA-containing hydrogel exhibited predominant cell fusion and cell chain formation within the 3D scaffold, a feature that remained less prominent even after 14 days in the 5G control sample. Image analysis quantified the number of cell chains containing three or more nuclei over time **(Figure 7C)**. At days 3, 7, and 14, myotube length was significantly higher in the TMA-containing sample compared to the control GelMA sample. For example, after 14 days, the average myotube length was 266.84 ± 52.24 µm in 5G and 358.19 ± 101.2 µm in 5G10TMA, demonstrating a significant difference between the groups. Similarly, the average number of cells per myotube **(Figure 7C)** showed a marked increase in the TMA group. Over the culture period, the cell count per myotube increased from 4.2 ± 1.4 to 8.8 ± 2.6 in 5G and from 10.1 ± 3.3 to 12.9 ± 4.5 in 5G10TMA, indicating that TMA promotes cell fusion within the scaffold even in couple initial days of culture. Additionally, cell morphology after an extended culture period (28 days) for all TMA and GelMA combinations, along with image analysis, is shown in **Figure S8**. A similar trend was observed for 5G1TMA and 5G5TMA, with higher TMA concentrations leading to enhanced myoblast proliferation and fusion. Higher magnification images of Actin staining **(Figure S9)** provide a clearer visualization of cell fusion and myogenesis promotion. To investigate whether physically blended Taur in GelMA would have a similar effect as TMA, cell encapsulation experiments were performed using 5G, 5G1T, 5G5T, and 5G10T, with F-actin staining results shown in **Figure S10** and **Figure S11**. The findings indicate that Taur alone enhances cell proliferation, but the cells remained mostly singlet and did not fuse significantly even till 14 days of culture as shown in higher magnification images **(Figure S11**). As shown in previous sections, most of the blended Taur within the GelMA matrix was released into the media within the first few days of swelling, suggesting that Taur alone does not have a prolonged effect on C2C12 cells. After 14 days of culture, cell chains were observed in Taur-containing scaffolds, similar to the control sample, indicating that C2C12 cells were undergoing natural myogenesis as seen in the control group **(Figure S10A)**. However, during the first week, the presence of Taur within the scaffold enhanced cell proliferation in the Taur-containing samples compared to GelMA alone. This indicates a higher number of single cells present within the scaffold. However, the image analysis presented in **Figure S10B** shows lower values for both myotube length and cell count per myotube. Moreover, the myotube length across different GelMA–Taur combinations appears relatively similar after day 7, likely because most of the Taur had been released into the media by that time. On the other hand, the cell count per myotube is significantly higher in the Taur-containing samples, as early-stage proliferation was enhanced within the first few days of culture, resulting in more available cells attempting to fuse and form myotubes **(Figure S10)**. In contrast, in TMA-containing samples, cells tended to exit the proliferation cycle earlier and initiated fusion, leading to the formation of longer myotubes **(Figure S8)**. Additionally, higher magnification images confirmed that predominant cell connections and myotube formation became more pronounced after 21 days of culture **(Figure S11)**. This finding supports the prolonged retention ofTMA within the scaffold and its continued influence on myogenesis.

To assess myogenic differentiation, Myosin Heavy Chain (MHC) immunofluorescence staining—a key protein required for myotube formation—was performed over 14 days of culture. **Figure 7D** illustrates MHC expression in C2C12 myoblasts within 50 and 5010TMA scaffolds. In the 50 scaffold, MHC-positive cells were first observed after 14 days of culture. In contrast, in the 5010TMA sample, most cells were already MHC-positive by day 3, and significant myotube formation was evident after 7 days, further increasing by day 14, indicating tissue maturation within two weeks. Quantitatively, the MHC-positive index, representing the ratio of MHC-positive cells to total cells, was calculated. **Figure 7E** shows that by day 3, 98.5 ± 1.3% of cells in 5010TMA were MHC-positive, while in 50, this number remained below 2%. During the initial days of culture, C2C12 myoblasts exit the cell cycle and commit to differentiation. MHC expression begins before fusion, marking the early differentiation phase. These MHC-positive single cells remain mononucleated, indicating that they have initiated muscle-specific gene expression [77,78]. After day 3, nearly all 5010TMA cells were MHC-positive, whereas in the 50 control sample, the MHC-positive ratio increased only to 6.5 ± 0.55% and 49.78 ± 8.18% by days 7 and 14, respectively. Moreover, MHC-positive cells in both groups underwent fusion, forming multinucleated myotubes. Cell-cell adhesion proteins (e.g., N-cadherin, M-cadherin) and fusogenic factors (e.g., Myomaker, Myomerger) increased over time, facilitating membrane merging [79–81]. To quantify myotube formation, the fusion index—the ratio of MHC-positive cells with more than 2 nuclei to total MHC-positive cells—was calculated. **Figure 7F** presents the fusion index of 5G and 5G10TMA over 14 days of culture. By day 3, no myotubes were observed in 5G, while the fusion index in 5G10TMA was already 7.3 ± 2.6%. After 14 days, the fusion index increased to 22.24 ± 0.76% in 5G and 86.8 ± 1.82% in 5G10TMA, demonstrating the significant impact of TMA on myogenesis. Additionally, merged images ofF-actin and MHC immunostaining are presented in **Figure S13**, showing MHC fiber alignment with F-actin-stained cell chains (green). Other TMA concentrations were also examined via MHC staining to assess the effect ofTMA concentration on myogenesis. **Figure S14** displays F-actin and MHC immunostaining of 5G1TMA and 5G5TMA scaffolds over 14 days of culture. In the 5G1TMA sample, MHC expression began around day 7, and cell fusion was initiated by day 14. In contrast, in 5G5TMA, MHC expression was detected as early as day 3, and by day 7, most cells were MHC-positive, with significant myogenesis occurring by day 14.

Furthermore, 3D bioprinted, cell-laden 5G and 5G10TMA scaffolds were cultured for one week, and their MHC immunostaining results are shown in **Figure 7G**. In scaffolds ∼400 µm thick, myogenesis was evident within a week in 5G10TMA, whereas 5G showed minimal myotube formation. Additionally, after photo-crosslinking, some cells on the glass surface were intentionally left unwashed to allow growth on a 2D surface. The blue background in **Figure 7G** represents a confluent layer of proliferating C2C12 cells on the glass substrate outside the 3D gel structure. However, these 2D-cultured cells showed no MHC secretion, reinforcing that both 3D cell culture and the presence ofTMA are critical for promoting C2C12 myogenesis. Additionally, **Figure 7H** shows the 3D confocal imaging of the bioprinted samples, while **Video S2** illustrates the distribution of F-actin and MHC-positive cells within the 3D structure of the 5G10TMA sample.

## 4. Conclusion

In this work, Taur was successfully functionalized with methacrylate groups to synthesize TMA, enabling covalent bonding with the GelMA backbone. The resulting hydrogel system demonstrated improved crosslinking control and structural stability with suitable mechanical, electrical conductivity, and swelling properties. Compared to physically incorporated Taur, TMA showed prolonged retention in the GelMA network, attributed to covalent integration. The incorporation of TMA also enhanced 3D printability, allowing for the fabrication of high-resolution and structurally defined constructs. Significantly, the bioactivity introduced by Taur and TMA supported C2C12 proliferation and myogenic differentiation, with TMA-containing scaffolds promoting more pronounced MHC expression and myotube formation. Overall, GelMA-TMA bioinks offer a promising strategy for skeletal muscle tissue engineering, combining structural precision with biological functionality for advanced biofabrication applications.

## Abbreviations

3D: Three-dimensional
SMTE: Skeletal muscle tissue engineering
GelMA: Gelatin methacrylate
DLP: Digital light processing
Taur: Taurine
TMA: Taurine methacrylate
SEM: Scanning electron microscopy
GMA: Glycidyl methacrylate
LAP: Lithium pheny1 2,4,6 trimethyl
PBS: Phosphate buffered saline
MHC: Myosin heavy chain

## Acknowledgement

This work was supported by a Natural Sciences and Engineering Research Council of Canada (NSERC) Discovery Grant (RGPIN-2020-04559) and Canada Foundation for Innovation John R. Evans Leaders Opportunity Fund.

