## supplemental Files for "Bioprinting Taurine-Incorporated Gelatin Methacrylate Hydrogels for Enhanced Muscle Tissue Regeneration"

### Supporting Information

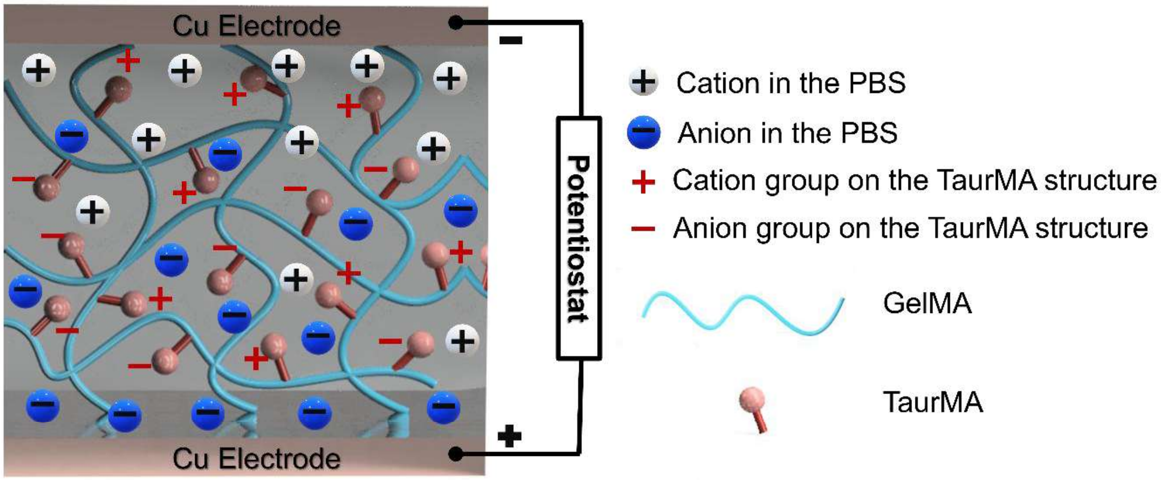


**Figure S1** *Schematic representation of the conductivity mechanism in GelMA hydrogels containing TMA.*

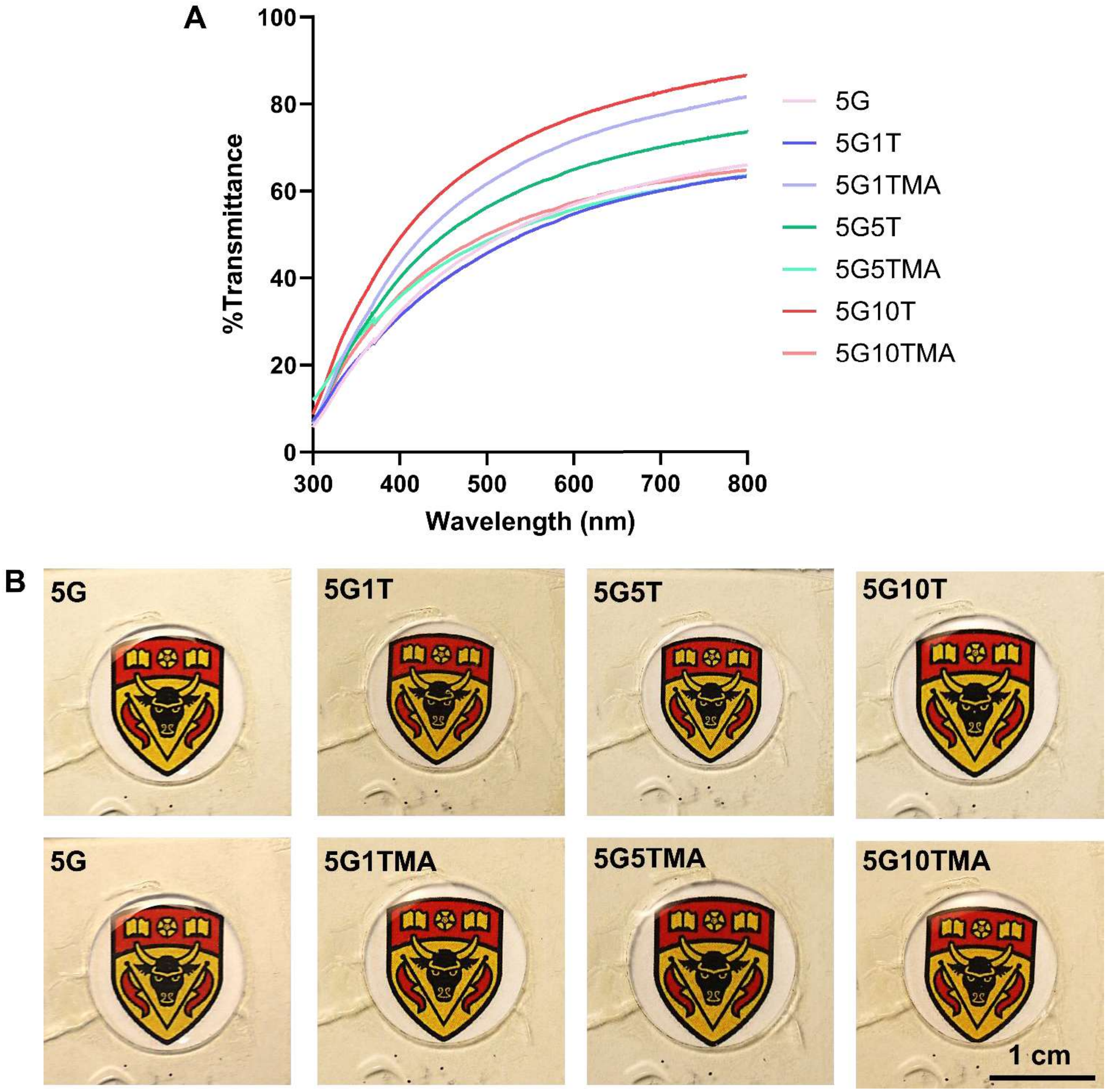


**Figure S2** *(A)Transmittance of the samples in the UV and visible light range and (B)Transparency evaluation of the prepolymer solution of GelMA containing Taur and TMA.*

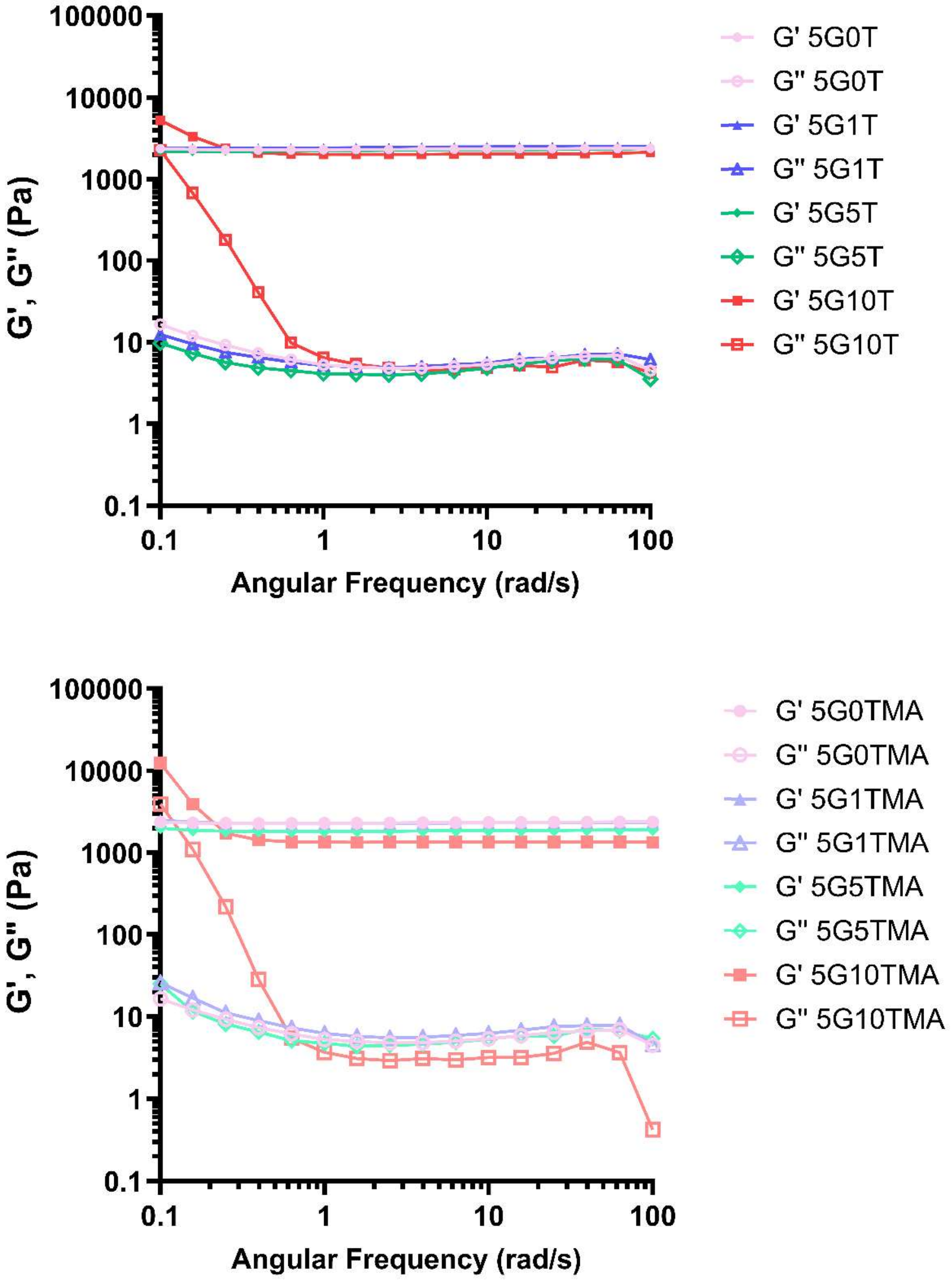


**Figure S3** *Angular frequency sweep of crosslinked GelMA hydrogels containing Taur and TMA.*

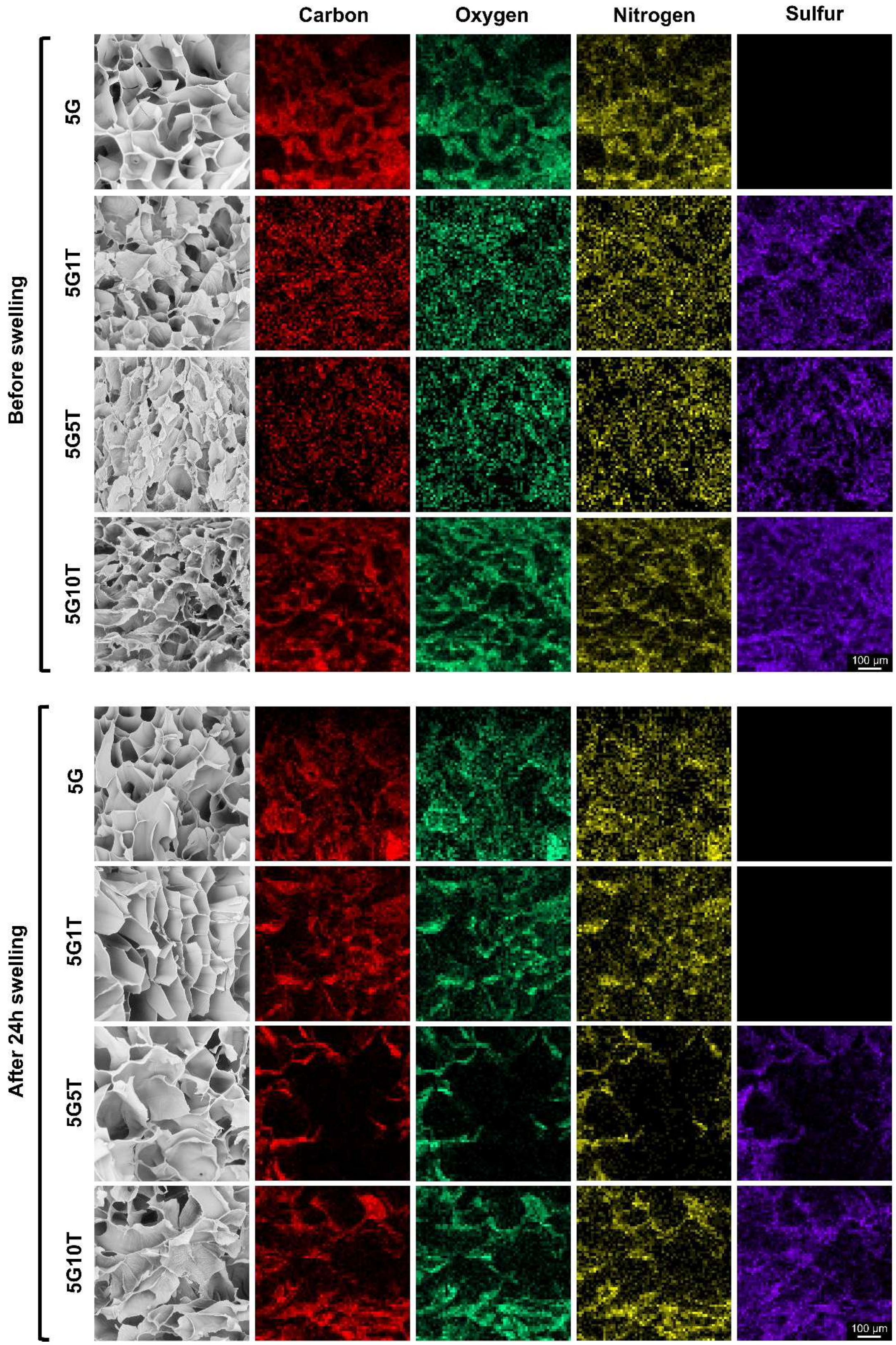


**Figure S4** *EDS Elemental Mapping of GelMA-Taur Hydrogels. SEM image overlaid with elemental distribution: red: Carbon, green: Oxygen, yellow: Nitrogen, and Purple: Sulfur, before and after 24 h swelling.*

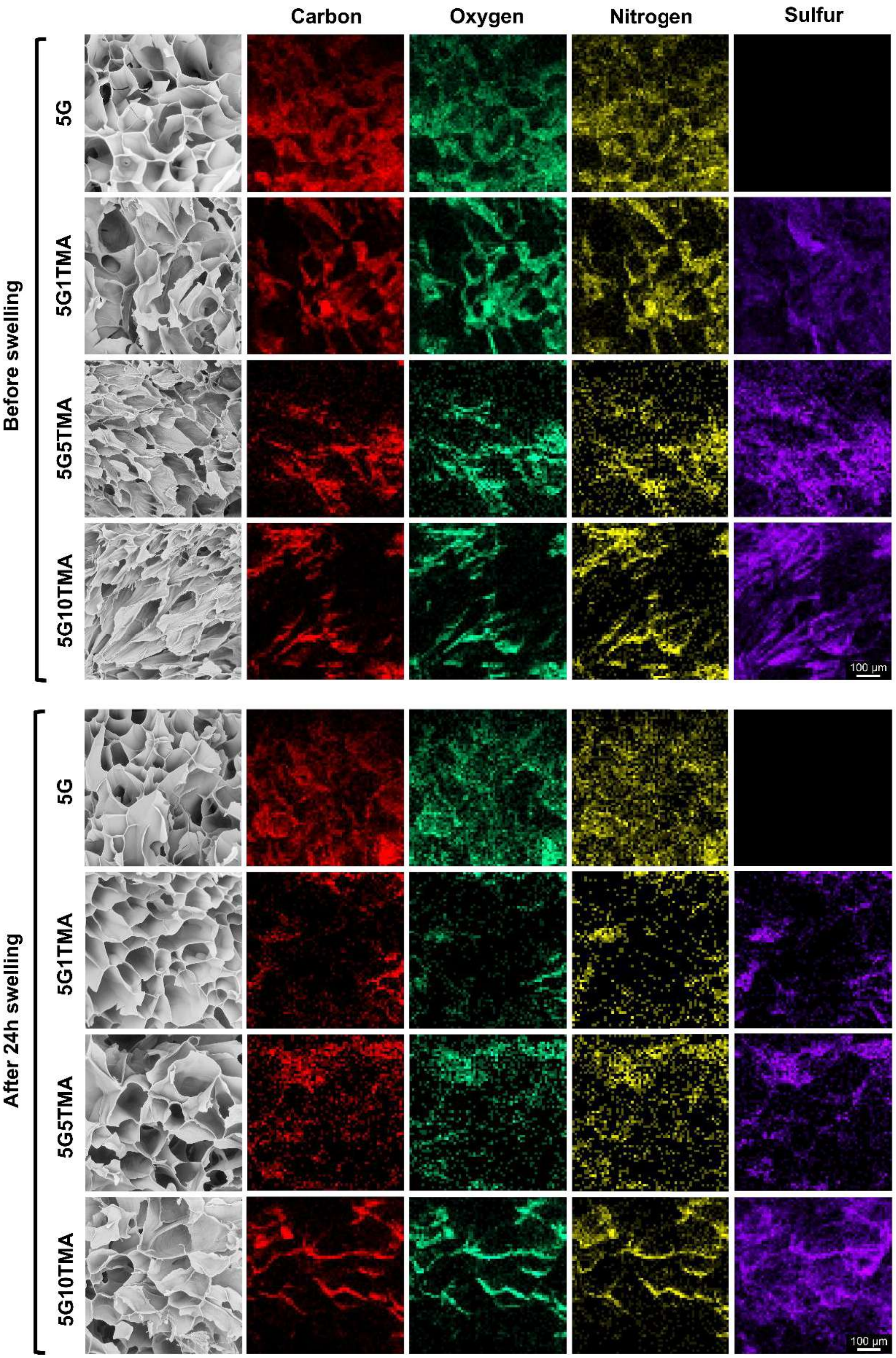


**Figure S5** *EDS Elemental Mapping of GelMA-TMA Hydrogels. SEM image overlaid with elemental distribution: red: Carbon, green: Oxygen, yellow: Nitrogen, and Purple: Sulfur, before and after 24 h swelling.*

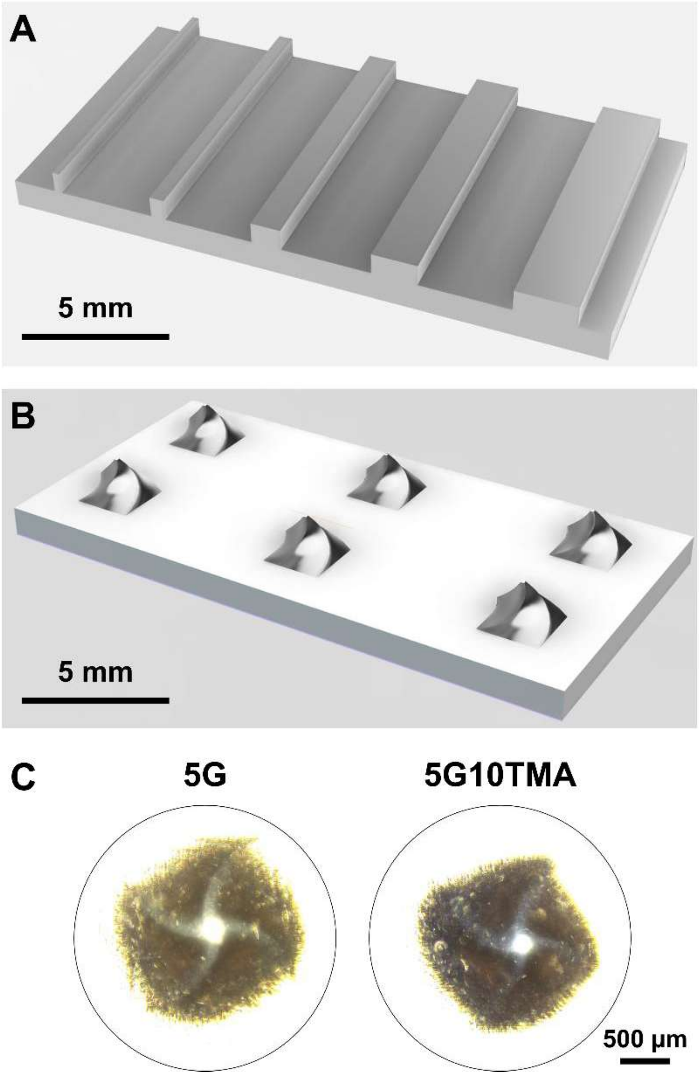


**Figure S6** *DLP 3D printing of 5G and 5G10TMA hydrogels. A) CAD model for evaluating the resolution of the printing. B) CAD model for twisted pyramid 3D printing. C) Brightfield imaging of printed twisted pyramids.*

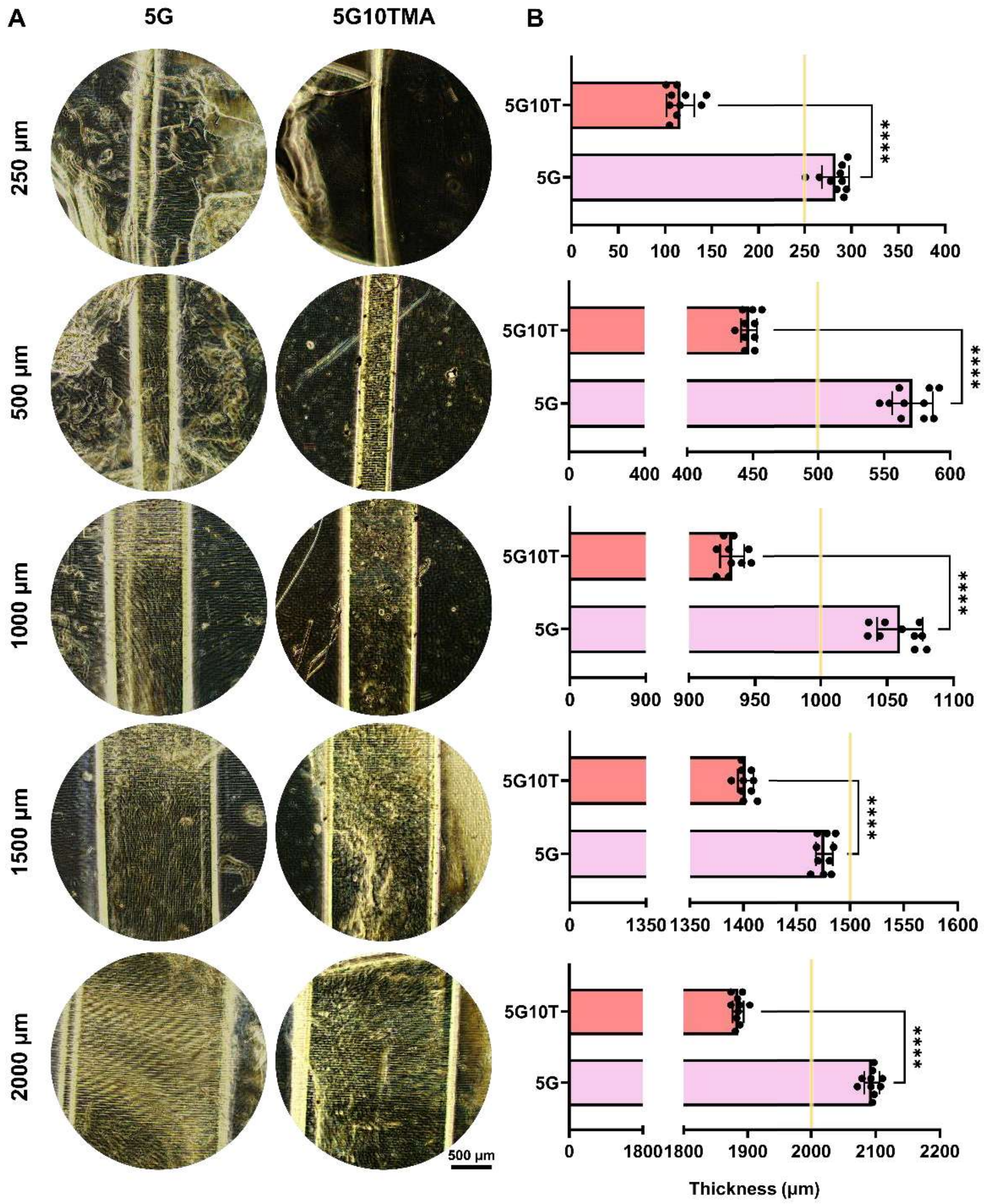


**Figure S7** *Resulution evaluation of DLP 3D printed structures using 5G and 5G10TMA hydrogels. A) Brightfield imading of line printing. B) Line thickness evaluation for different line printings.*

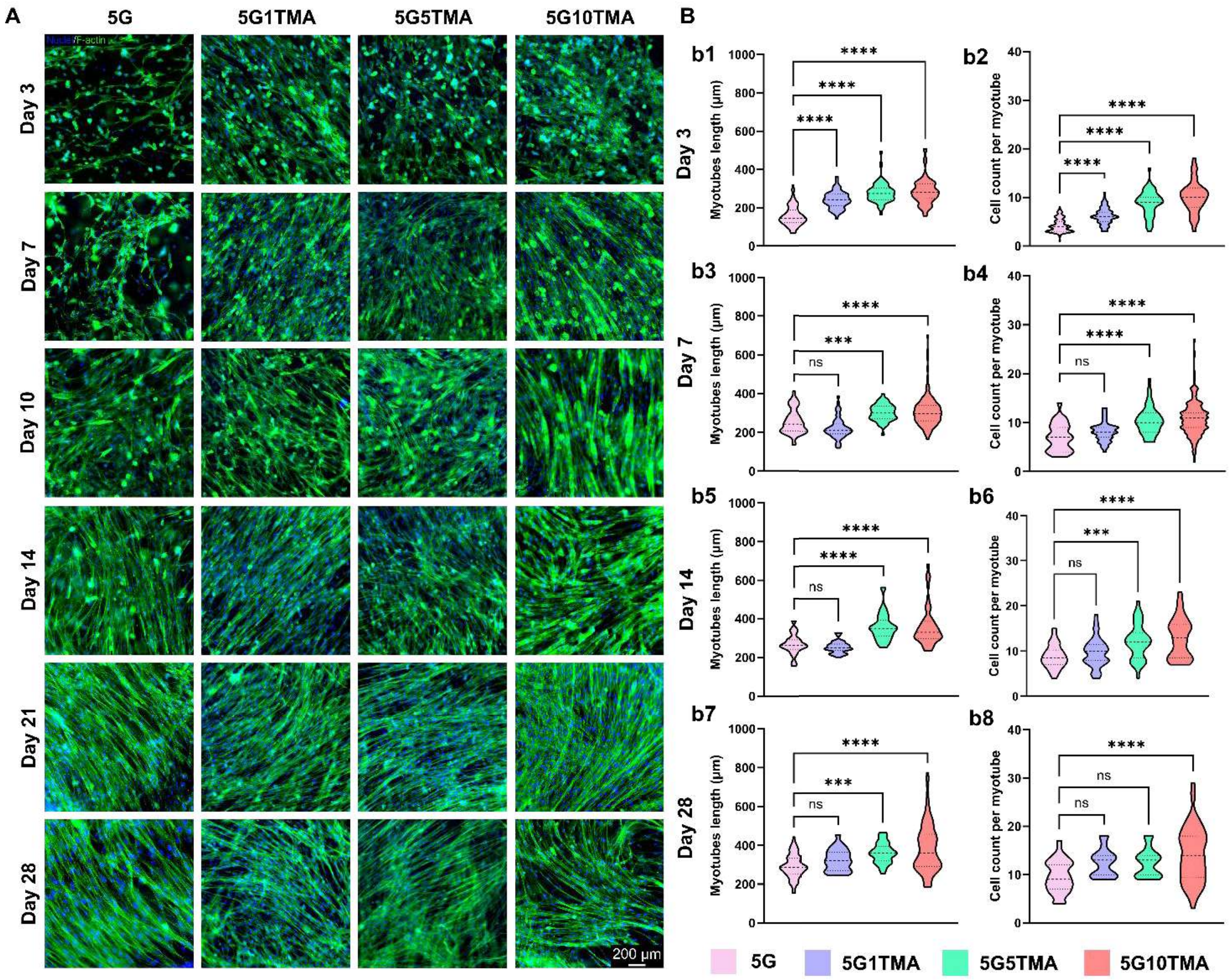


**Figure S8** *Morphological analysis of 3D-encapsulated C2C12 myoblasts. A) F-actin/DAPI immunostaining of GelMA containing 0, 1, 5, and 10% TMA cell-laden scaffolds for 28 days of culture. B) Image analysis evaluating myotube length and cell count per myotubes GelMA-based hydrogel cell-laden scaffolds at b1, b2) day 3, b3, b4) day 7, b5, b6) day 14 and b7, b8) day 28 of culture.*

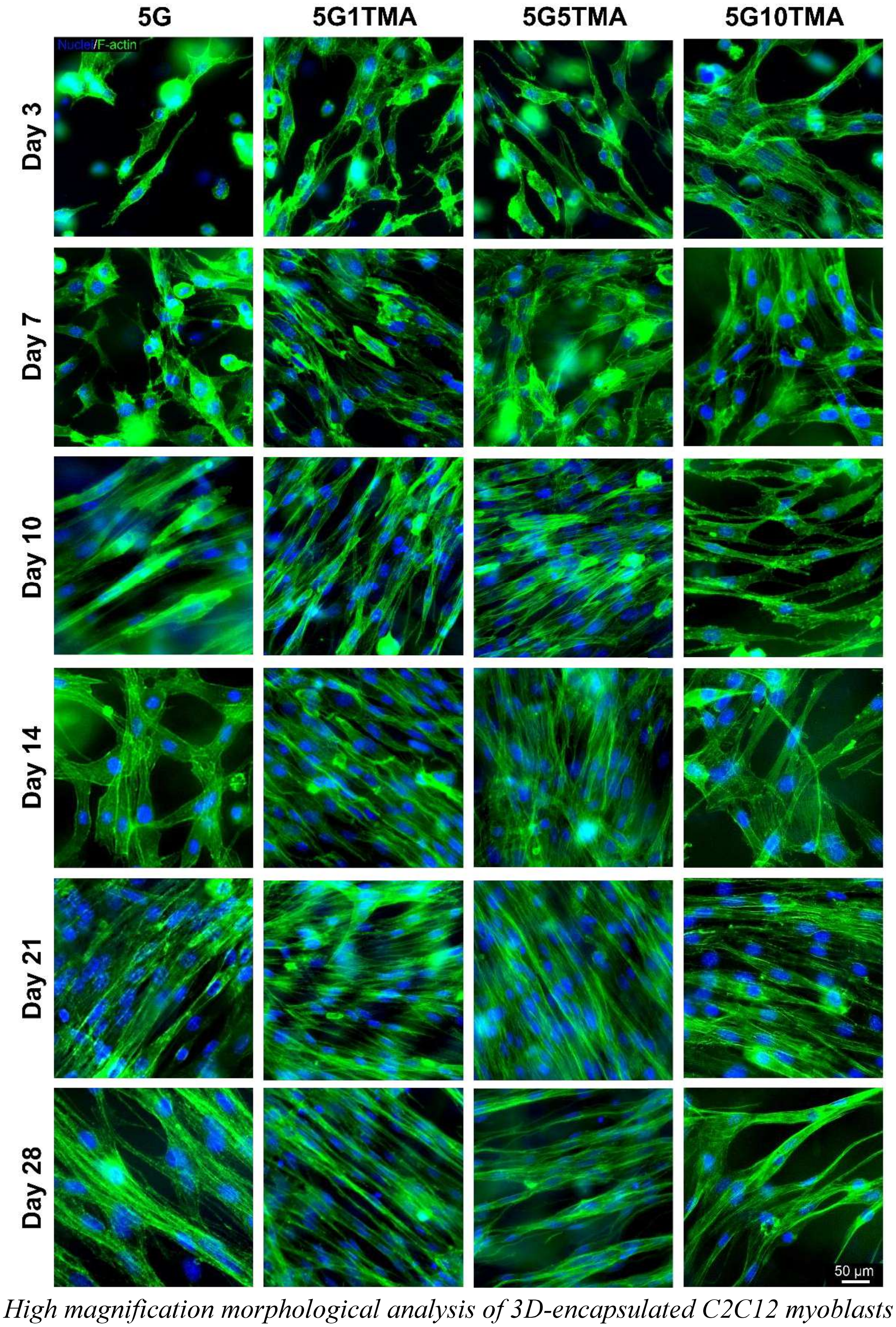


**Figure S9** *High magnification morphological analysis of 3D-encapsulated C2C12 myoblasts in GelMA hydrogels containing 0, 1, 5, and 10% TMA.*

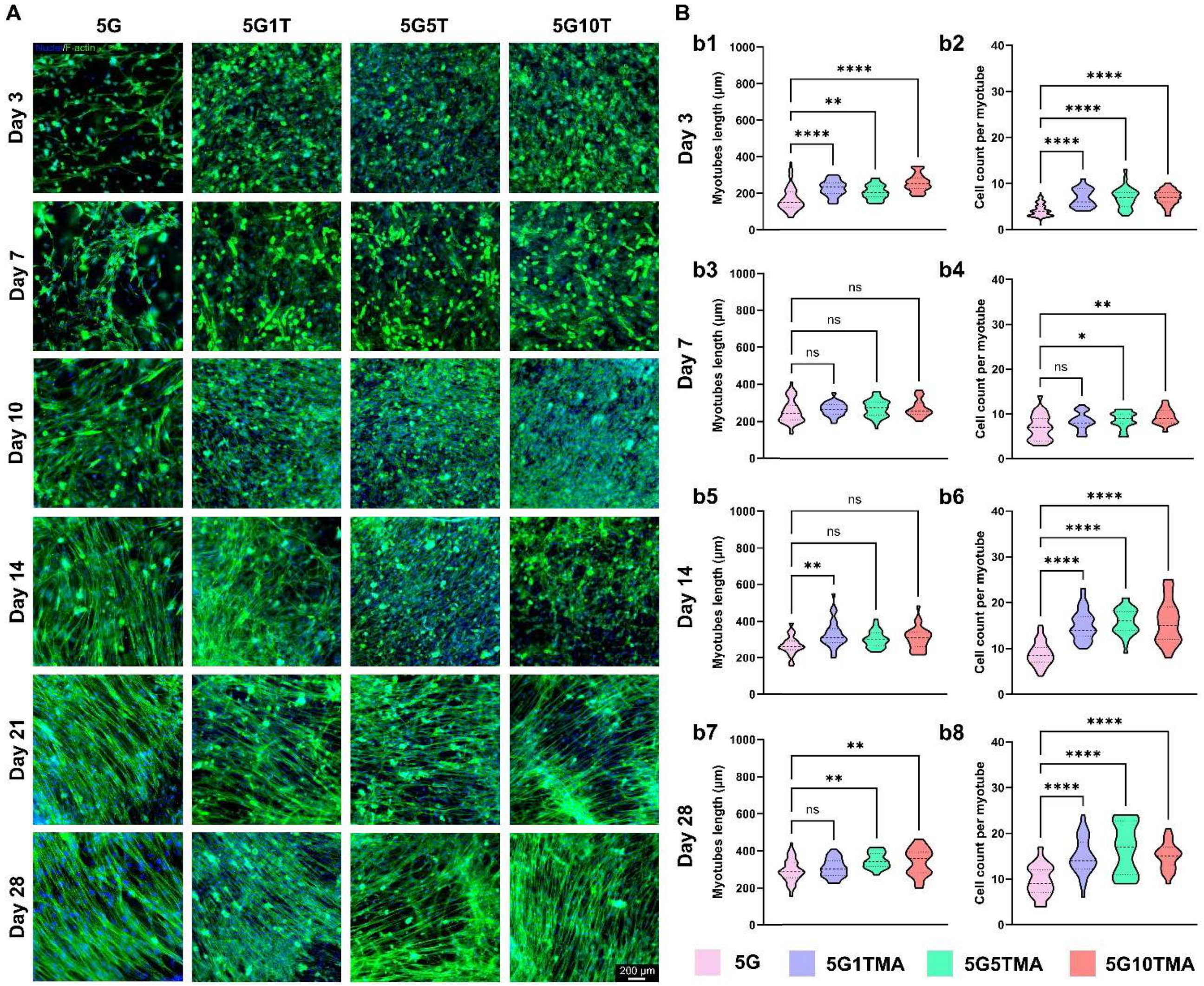


**Figure S10** *Morphological analysis of 3D-encapsulated C2C12 myoblasts. A) F-actin/DAPI immunostaining of GelMA containing 0, 1, 5, and 10% Taur cell-laden scaffolds for 28 days of culture. B) Image analysis evaluating myotube length and cell count per myotubes GelMA-based hydrogel cell-laden scaffolds at b1, b2) day 3, b3, b4) day 7, b5, b6) day 14 and b7, b8) day 28 of culture.*

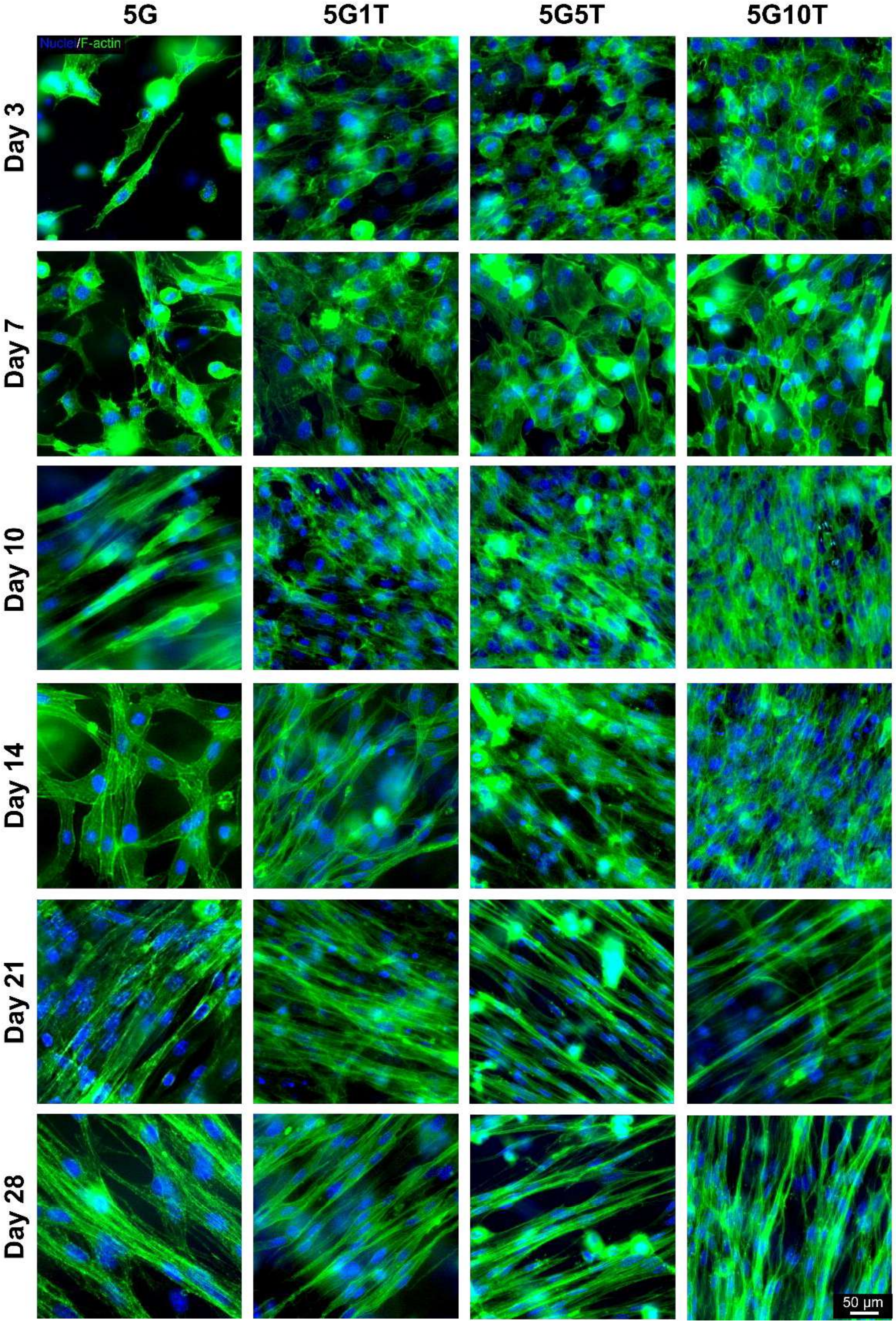


**Figure S11** *High magnification morphological analysis of 3D-encapsulated C2C12 myoblasts in GelMA hydrogels containing 0, 1, 5, and 10% TMA.*

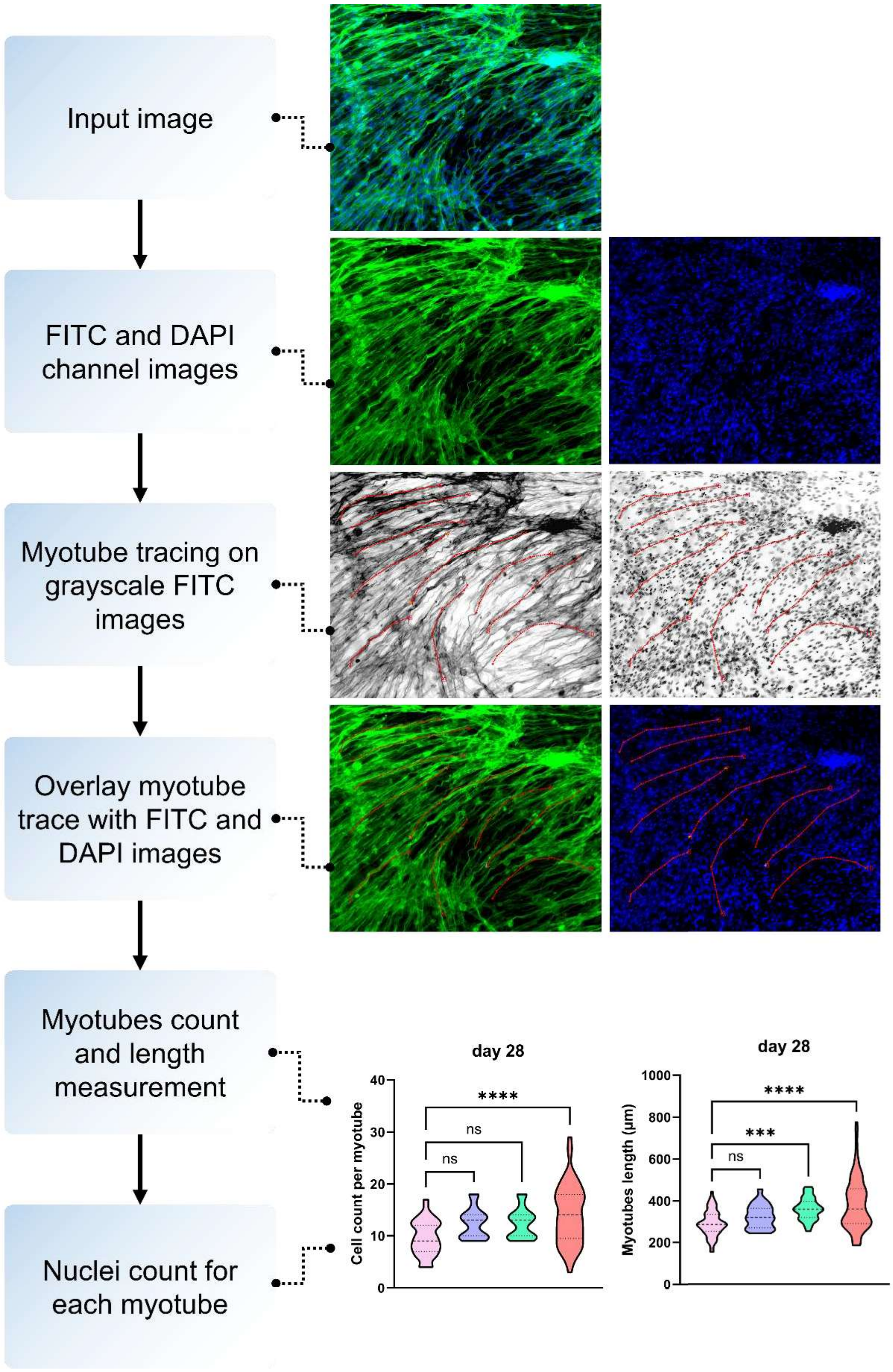


**Figure S12** *Image analysis process to evaluate myotube length and cell count per myotube using home-made MATLAB code.*

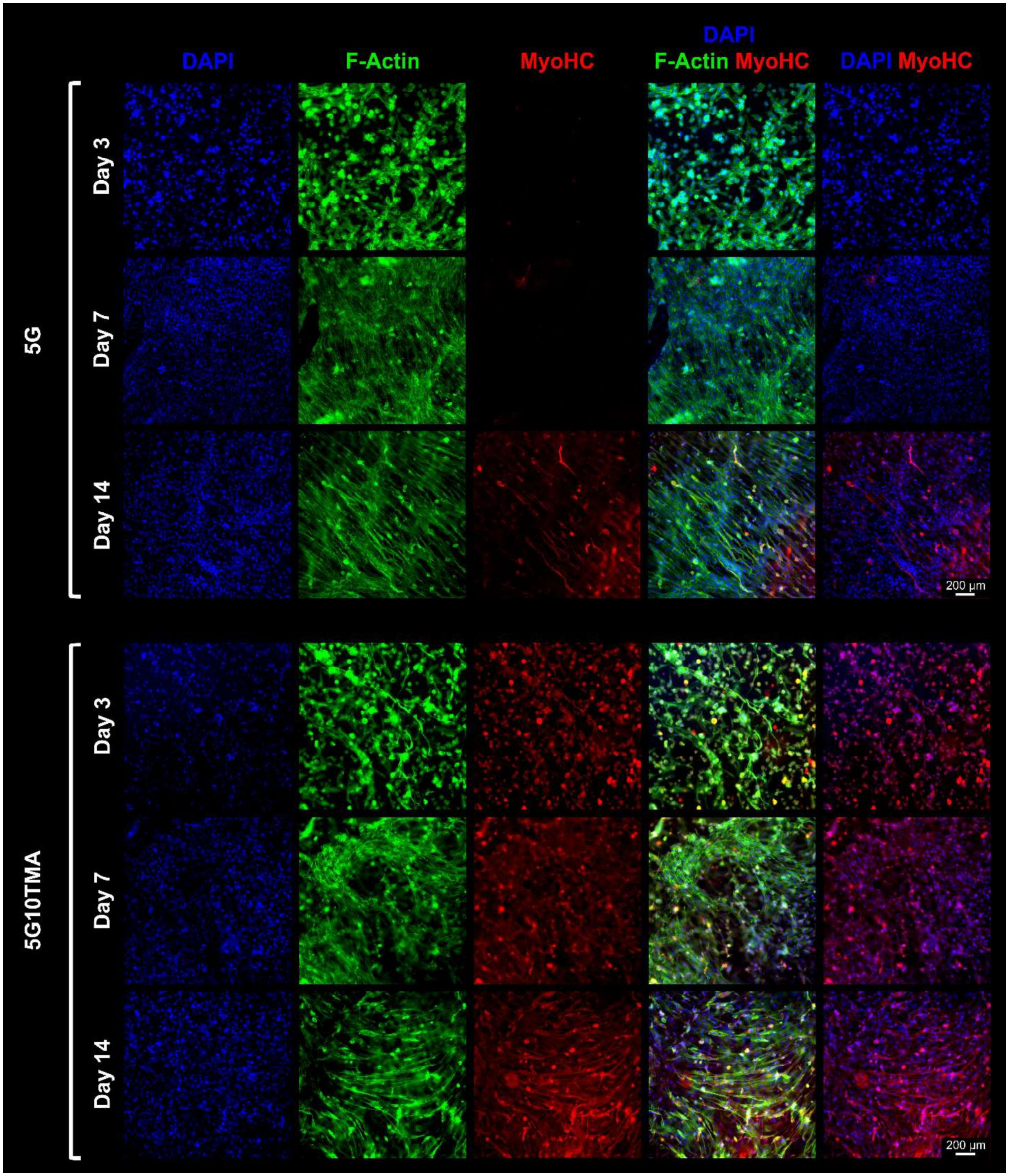


**Figure S13** *Morphological analysis and myogenic evaluation of 3D-encapsulated C2C12 myoblasts in 5G and 5G10TMA with F-actin/MHC/DAPI immunostaining.*

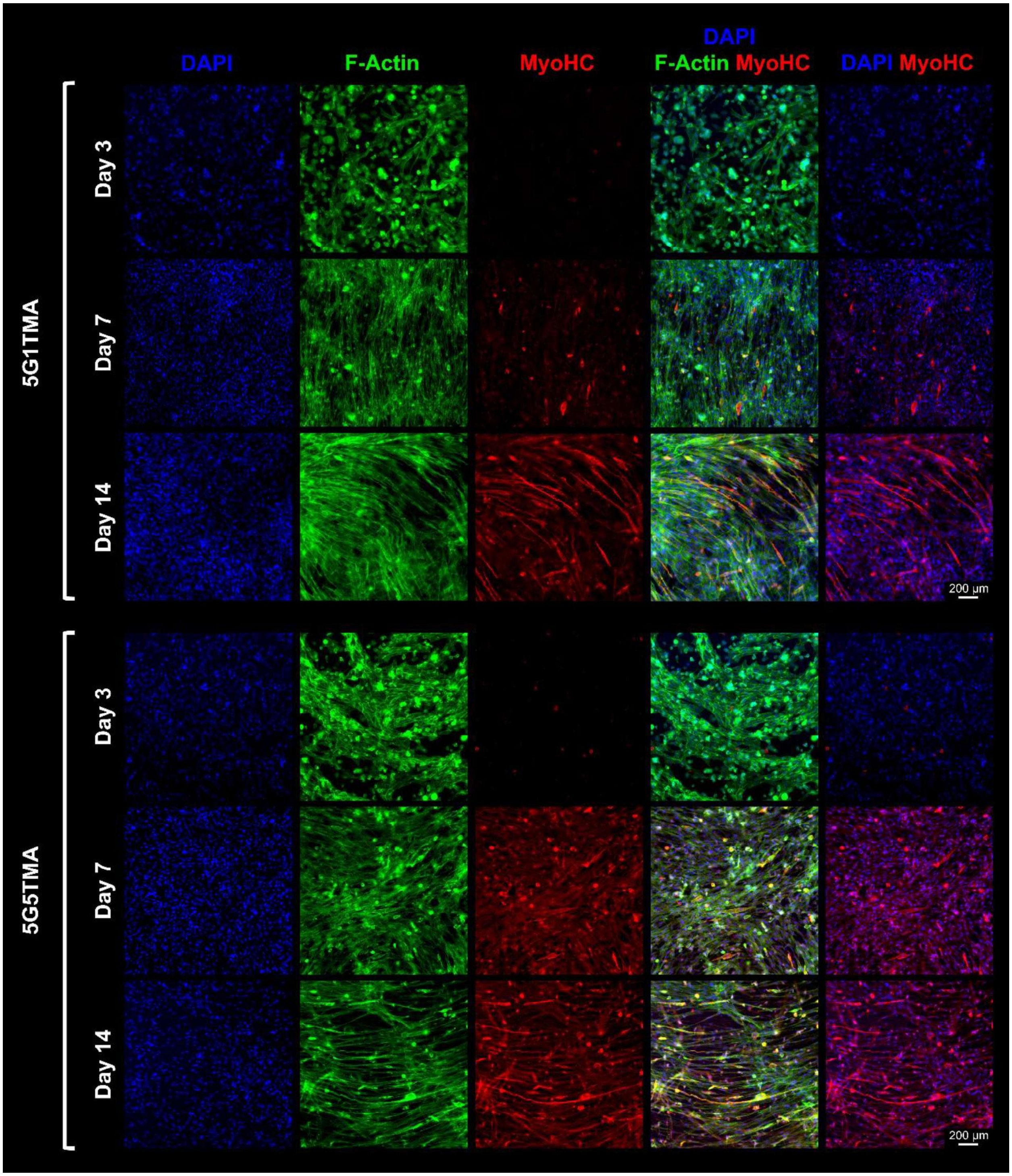


**Figure S14** *Morphological analysis and myogenic evaluation of 3D-encapsulated C2C12 myoblasts in 5G1TMA and 5G5TMA with F-actin/MHC/DAPI immunostaining.*

**Table S1** *Statistical analysis for hydrogels compression modulus*

| Tukey's multiple comparisons test | Summary | Adjusted P Value |
|---|---|---|
| 5G1T vs. 5G5T | ns | >0.9999 |
| 5G1T vs. 5G10T | ns | 0.9991 |
| 5G5T vs. 5G10T | ns | 0.9991 |
| 5G1TMA vs. 5G5TMA | ns | 0.6571 |
| 5G1TMA vs. 5G10TMA | * | 0.0485 |
| 5G5TMA vs. 5G10TMA | ns | >0.9999 |

**Table S2** *Quantitative elemental analysis for GelMA-Taur samples*

| | 5G | | 5G1T | | 5G5T | | 5G10T | |
|---|---|---|---|---|---|---|---|---|
| | Before | After | Before | After | Before | After | Before | After |
| C | 37.94 | 37.69 | 28.48 | 36.11 | 24.1 | 37.91 | 36.71 | 35.57 |
| O | 32.89 | 32 | 11.01 | 33.88 | 19.99 | 32.13 | 32.83 | 33.62 |
| N | 29.16 | 30.31 | 58.02 | 30.02 | 51.85 | 27.04 | 25.17 | 27.49 |
| S | 0 | 0 | 2.43 | 0 | 4.06 | 2.93 | 5.29 | 3.32 |

**Table S3** *Quantitative elemental analysis for GelMA-TMA samples*

| | 5G | | 5G1TMA | | 5G5TMA | | 5G10TMA | |
|---|---|---|---|---|---|---|---|---|
| | Before | After | Before | After | Before | After | Before | After |
| C | 37.94 | 37.69 | 37.46 | 30.75 | 34.5 | 19.59 | 35.77 | 35.7 |
| O | 32.89 | 32 | 33.68 | 35.28 | 29.48 | 34.08 | 31.2 | 32.2 |
| N | 29.16 | 30.31 | 25.83 | 33.69 | 27.96 | 41.04 | 22.2 | 24.5 |
| S | 0 | 0 | 3.02 | 0.28 | 8.15 | 5.29 | 10.82 | 7.6 |

**Video S1** *Twisted Pyramids 3D printing*

**Video S2** *Confocal microscopy*
